# NCI-Profiler: automated protein–ligand interaction detection and visualization

**DOI:** 10.64898/2026.10.11.758375

**Authors:** Sharma Rahul, Eun Yeol Lee

**Author notes:** Corresponding author: E.Y. Lee.

## Abstract

Non-covalent interactions govern ligand binding specificity and affinity, but existing analysis tools cover a limited set of interaction types, treat metal coordination as a generic contact, and struggle to depict macrocyclic ligands such as heme or cobalamin, leaving comprehensive interaction analysis fragmented across multiple programs. We developed Non-Covalent Interaction Profiler (NCI-Profiler), a free web-based and command-line tool for automated detection and visualization of nine types of protein–ligand non-covalent interactions, including metal coordination and halogen–water–hydrogen (XWH) bridges. NCI-Profiler detects interactions directly from PDB or mmCIF coordinates using literature-derived geometric criteria, without manual structure preparation, and produces interactive 2D diagrams, an in-browser three-dimensional viewer, and downloadable 3D PyMOL sessions, together with structured output files. Its accuracy was validated against 85 literature-annotated complexes and directly compared against PLIP, an established interaction-detection tool, on the same dataset. NCI-Profiler achieved F1-scores of 92–100% for π–π stacking, metal coordination, and water bridges, and 77–83% for the remaining types except hydrophobic contacts (54.8%); it detected literature-confirmed interactions more completely than PLIP in five of nine categories and ran roughly twice as fast. Three case studies further show its expanded coverage can reveal binding-pocket growth vectors, explain structure–activity trends, and distinguish receptor-subtype selectivity. NCI-Profiler is freely available at https://ncip.khu.ac.kr.

## 1. Introduction

Non-covalent interactions govern ligand binding specificity and affinity in protein–ligand complexes, making their accurate detection essential for drug design, enzyme engineering, and structural biology [1, 2]. The continued growth of structural databases and increasing complexity of protein–ligand assemblies have made manual characterization increasingly impractical, creating a need for automated approaches that can systematically identify and classify interactions from atomic coordinates. Over the past two decades, several tools have addressed this need. PLIP provides automated detection of seven interaction types from PDB files [3], while Arpeggio expands interaction coverage to fifteen subtypes using SMARTS-based atom typing [4]. PoseEdit provides interactive 2D diagrams through the ProteinsPlus server, whereas LigPlot+ generates static 2D interaction schematics [5, 6].

Despite these advances, protein–ligand interaction analysis remains distributed across tools with different interaction coverage, visualization capabilities, and output formats. Important interaction classes such as halogen–water–hydrogen (XWH) bridges are not reported by the web versions of commonly used profilers such as PLIP and Arpeggio, while metal coordination is generally treated as a contact rather than as part of a broader coordination analysis. Macrocyclic ligands such as heme, chlorophyll, and cobalamin also remain difficult to represent in conventional automated 2D interaction diagrams. In addition, workflows that combine comprehensive interaction detection with editable 2D visualization, 3D structural representation, and machine-readable output often require multiple programs. These limitations are particularly relevant when the objective is to obtain a complete and interpretable interaction profile directly from a protein–ligand structure.

We developed NCI-Profiler as an integrated web-based and command-line platform for automated protein–ligand interaction analysis. The platform detects nine classes of non-covalent interactions, including XWH bridges, and provides complementary structural, visual, and machine-readable representations of the resulting interaction profile. Its workflow accommodates structurally diverse complexes, including complexes containing multiple molecular components and macrocyclic ligands, while supporting both individual structure analysis and large-scale computational processing. The web server provides interactive analysis and visualization, whereas the command-line interface supports reproducible batch processing and programmatic integration into computational workflows.

We evaluated NCI-Profiler using 85 literature-annotated protein–ligand complexes spanning all nine interaction classes and quantified its agreement with reported interactions using precision, recall, and F1-score. This analysis showed F1-scores of 92–100% for π–π stacking, metal coordination, and water bridges, and 77–83% for the remaining interaction types. We then applied the workflow to three structural-analysis questions. In *S. cerevisiae* chitinase, water-mediated halogen contacts helped identify a fragment position associated with a subsequently validated binding pocket. For PDE5 inhibitors, the interaction profiles captured a shift from direct halogen bonding to water-mediated halogen contacts across a halogen series. In melatonin receptors, differences in the geometry of contacts around a conserved phenylalanine distinguished the binding modes of the same ligand across receptor subtypes. Together, these analyses evaluate NCI-Profiler across a progression from systematic interaction detection and classification to the interpretation of structure–activity relationships, binding-site expansion, and subtype-selective recognition.

## 2. Key Capabilities

NCI-Profiler combines interaction analysis with structure handling, visualization, and computational output in a single workflow. First, in terms of broad structural handling, the platform can analyze unprepared protein–ligand complexes and automatically distinguish relevant molecular components within structures containing multiple small molecules. This reduces the need for manual structure curation and supports consistent analysis across heterogeneous structural datasets. Second, for extended interaction profiling, in addition to commonly analyzed non-covalent interactions, NCI-Profiler explicitly identifies XWH bridges as a separate interaction class, capturing halogen-water-mediated contacts that would otherwise be missed if halogen interactions and water bridges were considered independently, and providing additional interpretive value for halogenated ligands. Metal-mediated interactions are also incorporated into the interaction profile, allowing these contacts to be considered together with other protein–ligand interactions. Third, for flexible structural visualization, interaction profiles can be examined through linked 2D and 3D representations. The editable 2D representation supports complex binding sites and macrocyclic ligands, while the three-dimensional context can be explored directly in the browser through an interactive viewer or through a downloadable PyMOL session. Fourth, for multi-ligand representation, multiple ligand interaction profiles from the same structure can be retained within a common analysis and represented together in a single PyMOL session. This facilitates examination of interactions across interconnected ligand-binding sites. Finally, for computational accessibility, the same analysis framework can be used for individual structures or large collections through the web server, CLI, and Python API. Structured output allows interaction information to be retained for subsequent computational analysis and feature extraction.

## 3. NCI-Profiler Usage

NCI-Profiler is accessible through two interfaces that share the same interaction detection engine, geometric criteria, and interaction reduction procedure. The web server is designed for interactive structure analysis and visualization, whereas the command-line interface (CLI) supports scriptable and batch processing. The two interfaces differ in input and visualization options: the web server accepts PDB and mmCIF files, whereas the CLI currently accepts PDB files only.

### 3.1 Web-server, input and output

The user provides a protein–ligand complex as a PDB or mmCIF coordinate file, either by direct upload or by retrieving a structure from the RCSB Protein Data Bank using its four-letter accession code. No manual preprocessing, protonation, or format conversion is required. For mmCIF input, the web server internally converts the structure to PDB format before analysis. Structures containing multiple molecular components are presented through a selection interface that categorizes detected components and allows individual molecules to be selected for analysis. The interaction reduction filter is enabled by default but can be disabled when the complete interaction set is required. Distance and angle thresholds can also be adjusted through the parameter control panel before analysis (Figure 1A). Default thresholds are listed in Table S1.

**Figure 1.**
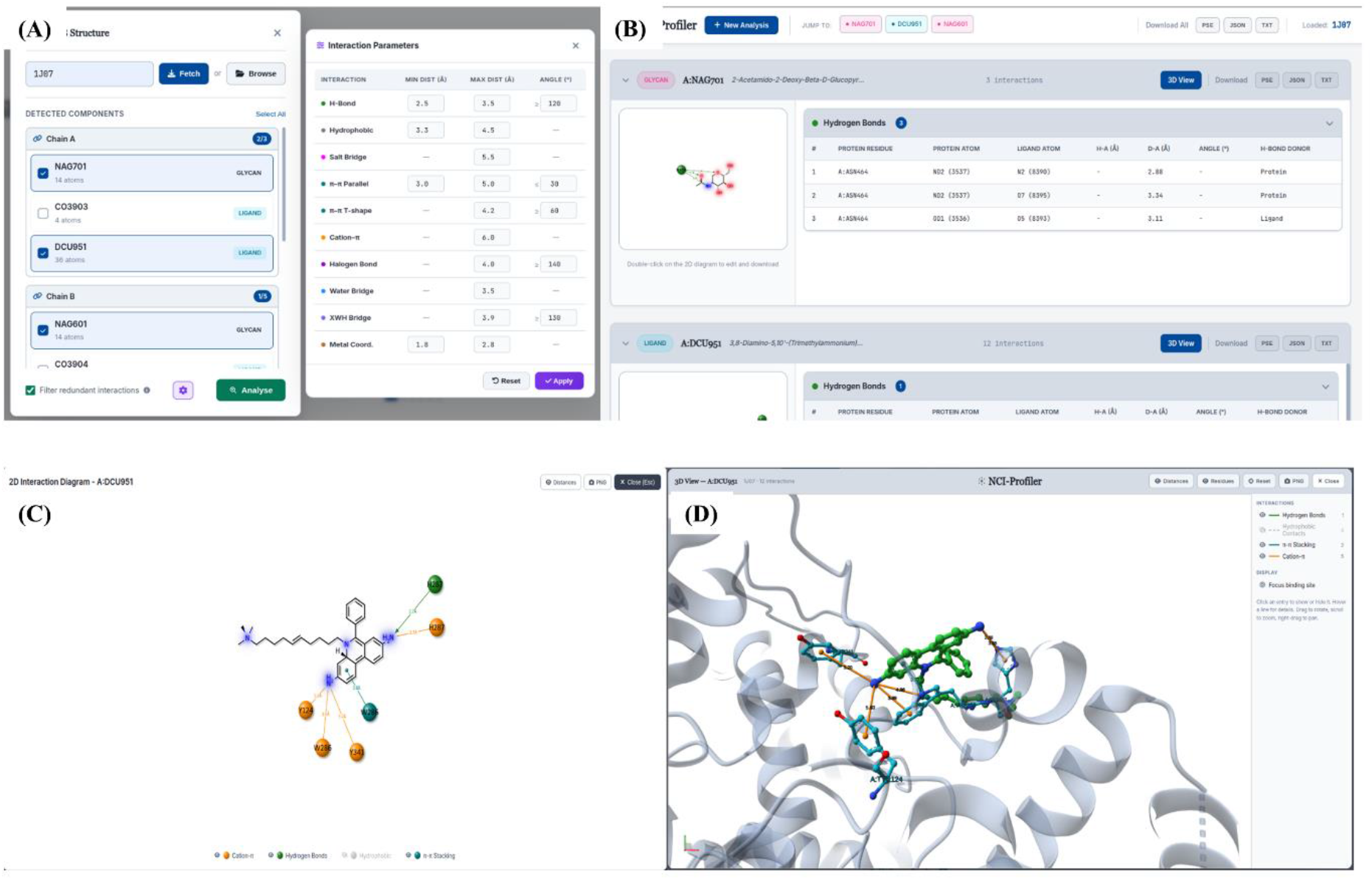
The NCI-Profiler web-server workflow. (A) Structure loading and multi-ligand selection interface for PDB entry 1J07, showing detected molecular components with individual selection checkboxes alongside the interaction-parameter panel for adjusting detection thresholds and angles. (B) Results page for the same analysis, showing per-ligand interaction tables, a 2D diagram thumbnail, and export options, with dedicated 3D View and Download controls provided for each ligand. (C) Interactive 2D diagram editor opened from the result thumbnail, with an interaction-type legend and export toolbar. (D) Interactive 3D viewer, opened from the 3D View button, shown here the ligand and interacting residues are displayed with interaction lines colored and labeled consistently with the 2D diagram, alongside an interaction-type legend with individual show/hide toggles and view/export controls.

After analysis, results for all selected ligands are displayed on a single page. Users can switch between ligands without rerunning the analysis, and each ligand is presented as an individual result card containing the interaction count, interaction-specific tables, and a thumbnail of the corresponding 2D diagram, and a 3D View button for opening an interactive three-dimensional view of the same interactions. The tables report the interacting residue, protein and ligand atom identifiers, interatomic distance, and, where applicable, geometric angle. Results can be exported at both global and per-ligand levels. Global export produces a combined result for all selected ligands, including a single PyMOL session containing their corresponding interactions. Per-ligand export produces a separate result for an individual ligand. This provides two complementary views of multi-ligand structures: a combined representation for examining interconnected binding sites and ligand-specific representations for focused analysis (Figure 1B).

The interactive 2D diagram can be opened from the result thumbnail. Residue labels can be repositioned to reduce overlap, individual interaction types can be shown or hidden through the color-coded legend, and distance labels can be toggled independently. The diagram can also be zoomed and edited before export at 600 DPI (Figure 1C). The three-dimensional viewer, opened from the 3D View button, renders the analyzed structure and its detected interactions directly in the browser using the Mol* viewer [7], without requiring locally installed software (Supplementary Information, Section 6). Interaction lines are colored and styled consistently with the 2D diagram, each interaction type can be independently shown or hidden through a corresponding legend, and residue labels, distance labels, a focused view of the binding site, and high-resolution transparent background image export are also available (Figure 1D). For offline analysis or publication-figure preparation, a corresponding PyMOL session is also available for download, providing a 3D representation of the analyzed binding site. The ligand, interacting residues, and interaction types are organized as separate PyMOL objects, allowing them to be independently displayed, hidden, or modified for further analysis and figure preparation. The 2D and 3D representations use the same interaction color scheme, as illustrated in Figure 2 for an EGFR kinase complex (PDB: 1XKK).

**Figure 2.**
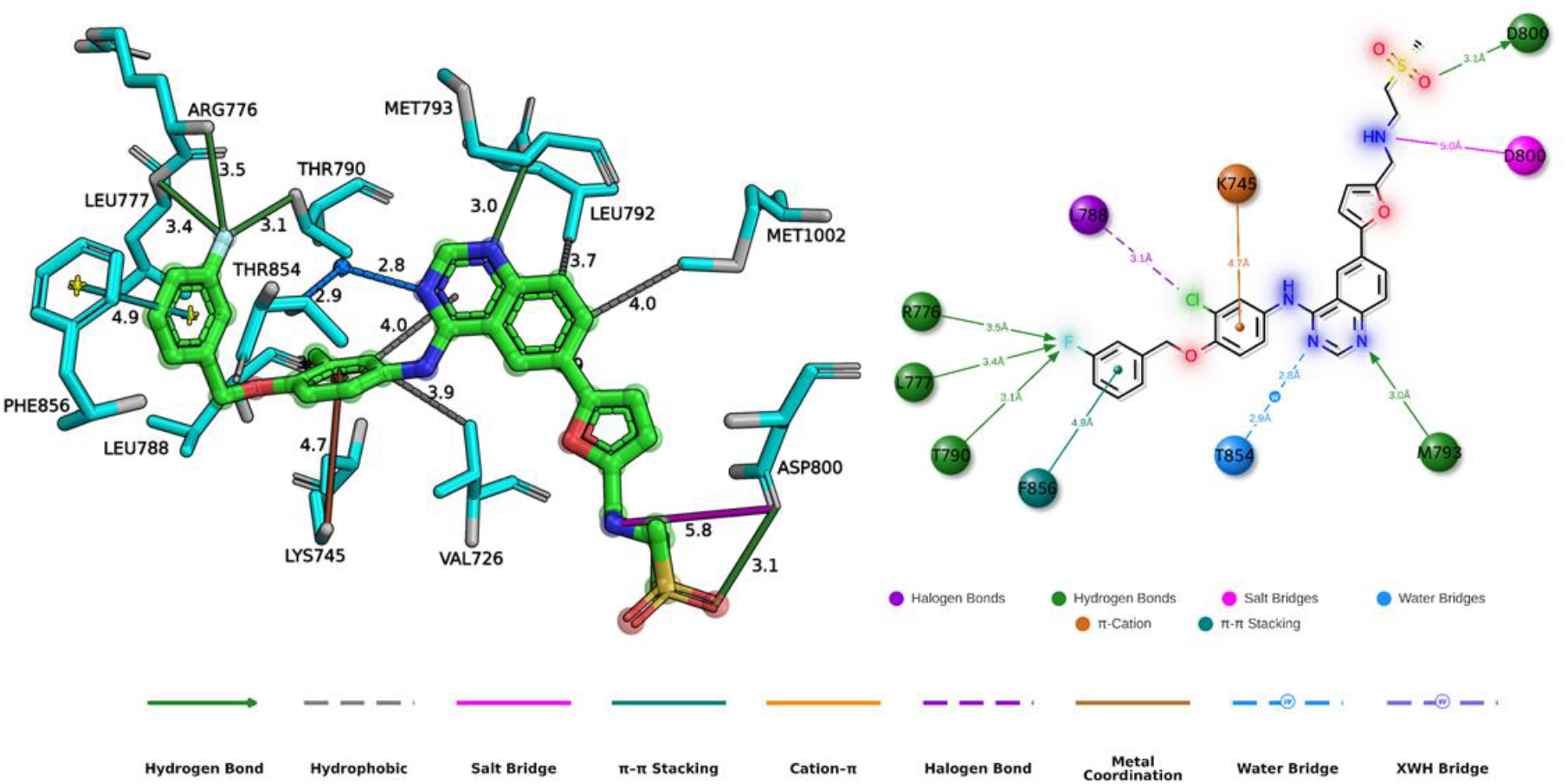
Representative NCI-Profiler output for EGFR kinase (PDB-1XKK, 2.40Å) with a quinazoline inhibitor (GW572016) showing complementary 3D (left, PyMOL session) and 2D (right, interactive diagram) visualizations of a protein–ligand binding site. Both representations use a unified color scheme for all nine interaction types, as defined in the bottom legend: hydrogen bonds, hydrophobic contacts, salt bridges, π–π stacking, cation–π, halogen bonds, metal coordination, water bridges, and XWH bridges. Residue spheres and interaction lines in the 2D diagram are color-coded by interaction type. Interatomic distances are labeled in Angstrom.

### 3.2 Command-Line Interface

The CLI is implemented in pure Python (>=3.8) with no mandatory third-party dependencies and is installable from the project repository (https://github.com/rahulsharma-bio/NCI-Profiler) using a standard “***pip install -e .***” command. Once installed, a single entry point, ncip, supports three modes of operation: (i) direct analysis of a local PDB coordinate file (***ncip structure.pdb***); (ii) on-demand fetching and analysis of RCSB entries by accession code (***ncip --fetch 1ABC***); and (iii) batch processing of entire directories of PDB files (***ncip --batch pdbs/ -o results/***). Ligand selection can be restricted by residue name (***-l***), chain identifier (***-c***), or residue number (***-r***), enabling precise targeting of a single binding site in multi-ligand or multi-chain structures. Results are written as: plain-text tabular report (***--txt***), structured JSON (***--json***), flat CSV (***--csv***), and a color-coded PyMOL script (***--pml***) using the same interaction color scheme as the web server.

The CLI exposes a Python API for programmatic integration, allowing the ***analyze()*** and ***print_results()*** functions to be imported directly and embedded in computational pipelines, molecular dynamics post-processing workflows, or virtual screening scripts. The CLI shares the same interaction detection engine, geometric criteria, and interaction reduction cascade as the web server. However, the CLI does not include the interactive 2D diagram, 3D PyMOL session file, or the real-time parameter adjustment interface, which are available exclusively through the web server at https://ncip.khu.ac.kr. Unlike the web server, the CLI currently accepts only PDB-format coordinate files; mmCIF input is not supported in the command-line release.

## 4. NCI-Profiler Algorithm

NCI-Profiler begins structure processing by classifying every molecular component in the input coordinate file into one of six categories (protein, solvent, common ion, cofactor, ligand, or glycan) using curated lookup tables; nucleic acid residues are excluded from analysis, and residues that do not match any curated entry default to the ligand category, ensuring that novel or chemically modified small molecules are captured without requiring user input. This classification engine was independently validated against the complete wwPDB Chemical Component Dictionary [8], achieving near-perfect accuracy of 99.996% across all categories (Supplementary Figure S1). Atomic charges required for salt bridge detection are inferred through a geometry-based approach that does not require explicit hydrogen atoms, eliminating the external protonation step required by tools such as PLIP [3] and Arpeggio [4]. In a benchmark of 100 structures analyzed individually under identical hardware conditions, NCI-Profiler was approximately 2.2-fold faster than PLIP on average, with a greater advantage for larger complexes. This difference becomes relevant for large structural databases, where per-structure overhead accumulates across thousands of complexes (Supplementary Information, Speed Benchmark, Supplementary Figure S10).

NCI-Profiler evaluates nine interaction classes against literature-derived geometric thresholds applied to every protein–ligand atom pair within the binding site (complete threshold values in Supplementary Table S1). Candidate atom pairs are first identified by retaining all protein, water, and metal atoms within a 12.0 Å radius of the ligand centroid before applying the type-specific distance and angle criteria for each interaction class. Halogen bonds and XWH bridges are evaluated using the σ-hole model [9], with XWH bridges additionally requiring a simultaneous hydrogen bond between the bridging water and a protein residue [10]. Metal coordination is assessed for a broad range of biologically relevant metal ions by comparing observed donor positions against reference coordination geometries, with the τ4 and τ5 indices computed to distinguish four- and five-coordinate geometries [11].

Raw geometric detection can assign the same atom to multiple interaction types simultaneously, for example when a charged donor atom qualifies as both a hydrogen bond and a salt bridge, producing overlapping and potentially misleading contacts in the output. NCI-Profiler resolves this redundancy through an eight-step specificity cascade applied after all nine interaction classes have been evaluated. The cascade prioritizes metal coordination first, then resolves overlap between salt bridges, halogen bonds, and hydrogen bonds, followed by ring-level and residue-level deduplication for the remaining interaction types (complete step-by-step logic in Supplementary Information).

For 2D diagram generation, the ligand is first assembled as a chemically correct molecular object with resolved bond orders using a three-tier cascade based on Open Babel [12] and RDKit (Supplementary Information, Section 5). Ligand coordinates are then generated by a competitive evaluation of two independent layout algorithms, CoordGen and RDKit’s rdDepictor, selecting the layout with fewer bond crossings. Macrocyclic ligands are handled separately using a CCD-template-based strategy via PDBe CCDUtils [13] when the ligand’s chemical component has a corresponding entry in the wwPDB Chemical Component Dictionary; otherwise, a PCA-based coordinate-flattening fallback is followed (Supplementary Information, Section 5). Interacting residues are then placed around the ligand using an automated angular-sector assignment followed by iterative collision resolution to minimize label overlap (Supplementary Information, Section 5). The resulting diagram is rendered as an interactive SVG with type-specific line conventions for each interaction class, draggable residue labels, toggleable interaction-type layers, and toggleable distance labels, and can be exported directly in .png format from the browser without server-side processing.

## 5. Results and Validations

NCI-Profiler was validated against 85 protein–ligand complexes (85 unique PDB structures) spanning all nine interaction types and resolutions from 1.10 to 3.50 Å. Structures were selected from the published literature only when the associated crystallographic or biochemical study explicitly reported specific non-covalent interactions at the binding site involving a specific amino acid residue and, where available, the specific ligand atoms involved. This criterion provided an independent, literature-derived ground truth against which NCI-Profiler’s automated detection could be directly compared, rather than relying on an internally generated or assumed reference. Among the structures meeting this criterion, the final dataset was assembled to include all nine interaction types. As a result, the dataset contains 7–13 complexes per interaction type (Supplementary Table S3). A contact was scored as a true positive (TP) only when it was detected by NCI-Profiler and also reported in the reference publication. A contact was scored as a false positive (FP) when NCI-Profiler detected an interaction that was not reported in the corresponding literature, and as a false negative (FN) when an interaction was reported in the literature but not detected by NCI-Profiler. Because each published study typically reports only the interactions considered relevant to its specific analysis, rather than an exhaustive list of every geometrically valid contact, a false positive in this scheme does not necessarily indicate an incorrect detection. It may instead reflect a genuine interaction that the original publication simply did not discuss. Performance was assessed using precision, recall, and F1-score for each interaction type independently, with full per-structure TP/FP/FN counts provided in Supplementary Table S2 and aggregated per-type results in Supplementary Table S3.

Briefly, metal coordination, water bridges, and π–π stacking achieved the highest performance (F1 = 100%, 97.9%, and 92.7%, respectively), consistent with the highly specific geometric criteria that define these interaction types. Cation–π interactions (F1 = 82.8%), XWH bridges (81.0%), hydrogen bonds (78.6%), salt bridges (78.6%), and halogen bonds (77.8%) also showed strong overall agreement with the literature. Most false positives and false negatives in these categories arose from contacts near the geometric detection thresholds rather than from misclassification. Hydrophobic contacts yielded the lowest F1-score (54.8%), with precision and recall equally balanced, indicating a symmetric rate of over- and under-prediction rather than a directional bias. This lower performance reflects the absence of a universally accepted distance cutoff for hydrophobic contacts in the literature, which limits the consistency of the ground truth itself rather than indicating a systematic detection failure. Users for whom hydrophobic contacts are central to their analysis are encouraged to consult the raw, unfiltered interaction output and adjust detection thresholds via the web server’s parameter control panel or the CLI’s --config file. Detailed per-structure results are provided in Supplementary Table S2, along with representative 2D/3D visualizations for a subset of structures from each interaction category in Supplementary Figures S2–S9.

To benchmark NCI-Profiler against an existing tool, both NCI-Profiler and PLIP were evaluated on the same 85 complexes using the same criteria. Based on F1-score, NCI-Profiler achieved higher values in four interaction categories, PLIP in three, with two categories showing the same F1-score for both tools. Recall provided a different comparison: NCI-Profiler achieved higher recall in five categories, PLIP in one, with the remaining three tied. Because recall measures the fraction of literature-confirmed interactions detected by a tool, it provides a more direct measure of missed detections without penalizing contacts that were detected by the tool but not reported in the reference literature (Supplementary Table S4).

## 6. Exemplary cases

The utility of NCI-Profiler extends across a range of structural-analysis contexts and is not limited to identifying individual non-covalent interactions. Its outputs can be used to interpret binding features, compare interaction patterns, and characterize complex molecular environments within experimentally determined structures. The following cases illustrate this broader utility across different applications.

### 6.1 Fragment-Based Drug Design, Growth Vector Identification (2UY3)

Fragment-based drug design (FBDD) relies on identifying productive growth vectors from initial fragment hits to guide elaboration into potent lead compounds. NCI-Profiler can facilitate this process by revealing hidden interactions at fragment peripheries that may indicate exploitable binding hotspots. We applied NCI-Profiler to the crystal structure of 8-chlorotheophylline bound to *S. cerevisiae* chitinase ScCTS1 (PDB: 2UY3). 8-chlorotheophylline was identified as a fragment hit against ScCTS1 through screening of the Prestwick Chemical Library, with a measured inhibition constant of Ki = 600 µM [14]. NCI-Profiler detected two XWH bridges in which the chlorine atom interacts through HOH2303 with Tyr32 (3.0 Å) and Phe210 (2.9 Å) (Figure 3). These water-mediated halogen contacts connect the fragment’s C-8 position to a deep active-site pocket occupied by ordered water molecules. Notably, the original authors identified this pocket as a key elaboration site, stating that “inhibitors possessing moieties that could occupy this pocket and displace these water molecules would give additional favorable entropic contributions to binding.” This prediction was validated by kinetin (N⁶-furfuryladenine), in which the furfuryl group extends into this pocket and displaces the bridging waters, achieving a Ki of 3.2 µM, a ∼190-fold improvement in affinity [14]. Thus, the XWH bridges detected at the fragment–pocket interface identify the C-8 position as an interaction feature associated with a growth vector that was subsequently validated through fragment elaboration. This case illustrates how NCI-Profiler can support structural interpretation in FBDD by linking hidden interactions to experimentally validated binding opportunities.

**Figure 3.**
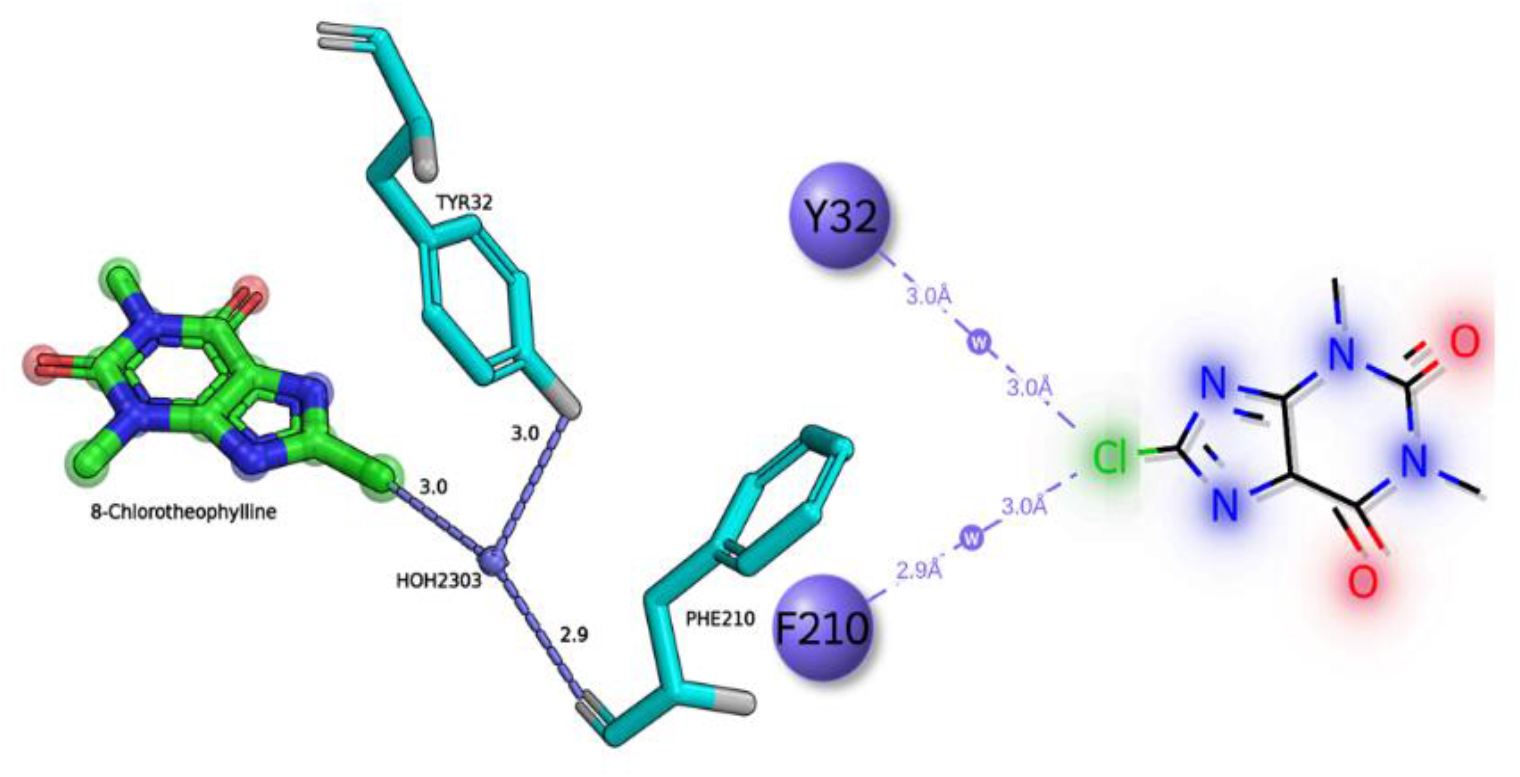
NCI-Profiler identifies a fragment growth vector through XWH bridge detection. Analysis of 8-chlorotheophylline bound to *S. cerevisiae* chitinase ScCTS1 (PDB: 2UY3). (Left) Three-dimensional view showing two XWH bridges (purple dashed lines) from the chlorine atom of 8-chlorotheophylline (green sticks) through bridging water HOH2303 (blue sphere) to Tyr32 and Phe210 (cyan sticks). Distances are labeled in Å. (Right) NCI-Profiler 2D interaction diagram depicting the same XWH bridges (purple dashed lines; w, water-mediated) connecting the Cl atom to Tyr32 (3.0 Å) and Phe210 (2.9 Å). These XWH bridges mark a growth vector toward a deep pocket; elaboration to kinetin, whose furfuryl group displaces these bridging waters, improves affinity 190-fold (Ki from 600 µM to 3.2 µM).

### 6.2 Halogen scan SAR analysis, PDE5 inhibitor series (3SHZ / 3SIE / 4OEW)

Structure–activity relationship (SAR) analysis relies on linking chemical modifications within a compound series to changes in molecular interactions and binding potency. NCI-Profiler can support this analysis by resolving interaction changes that arise from specific structural modifications, including interactions mediated through bridging molecules such as water. We applied NCI-Profiler to three PDE5 inhibitors from a systematic halogen scan series in which only the halogen at the pyrimidinyl-5C position differs [15, 16]. The chloro analog (3SHZ, IC₅₀ = 35.9 nM) shows no halogen-mediated contact: the Cl···O(Tyr612) distance of 3.60 Å exceeds the chlorine σ-hole cutoff, and only hydrogen bonds to Gln817 and π-stacking with Phe820 are detected (Figure 4, A). The bromo analog (3SIE, IC₅₀ = 13.3 nM) forms a direct halogen bond to Tyr612 (3.32 Å), consistent with its 2.7-fold potency gain (Figure 4, B). The iodo analog (4OEW, IC₅₀ = 7.2 nM) does not achieve a direct halogen bond (I···O = 3.68 Å). Instead, NCI-Profiler detects three XWH bridges in which the iodine engages a structured water molecule (HOH1005) that relays contacts to Tyr612, Asp764, and Ala767, extending the interaction network beyond the single residue contacted by the bromo analog (Figure 4, C). This water-mediated interaction network, which would not be captured by conventional interaction profilers that lack XWH detection, provides a structural explanation for the highest potency of the iodo derivative despite the absence of a direct halogen bond and identifies Asp764 and Ala767 as secondary contact points accessible through halogen–water relays. This case illustrates how NCI-Profiler can support SAR interpretation by resolving changes in interaction geometry and revealing water-mediated contacts that help explain potency differences across a chemical series.

**Figure 4.**
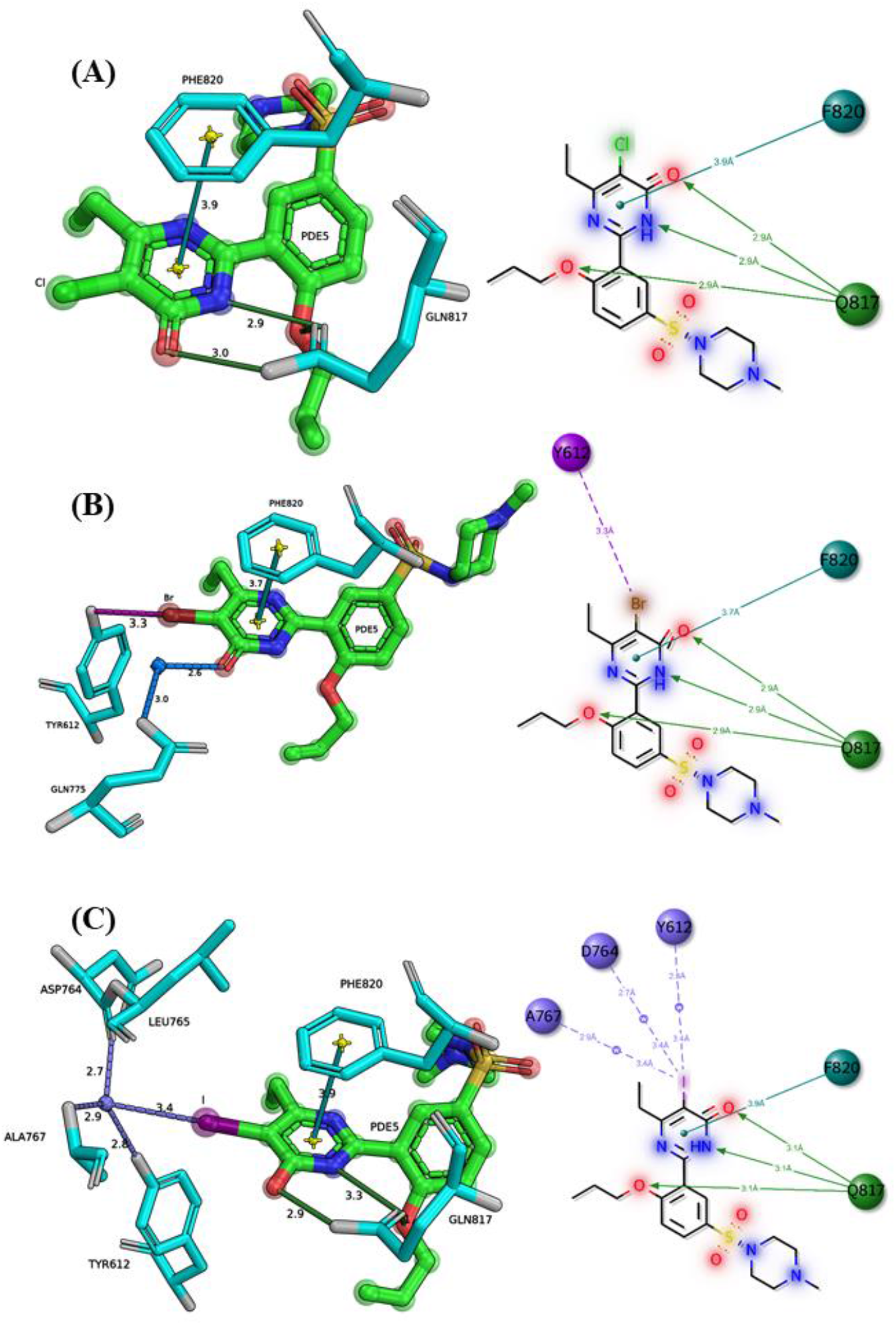
Halogen scan SAR analysis of PDE5 inhibitors. NCI-Profiler interaction profiles for three PDE5 inhibitors differing only in the halogen at the pyrimidinyl-5C position. (A) Chloro analog (3SHZ, 5CO): no halogen bond detected; only hydrogen bonds (green) to Gln817 and π-stacking (cyan) with Phe820 are present. (B) Bromo analog (3SIE, 5BO): a direct halogen bond (magenta, 3.3 Å) to Tyr612 is detected alongside hydrogen bonds and a water bridge (blue) to Gln775. (C) Iodo analog (4OEW, 5IO): three XWH bridges (purple) relay the iodine σ-hole through HOH1005 to Tyr612, Asp764, and Ala767. The interaction profiles correlate with the IC₅₀ rank order (I 7.2nM > Br 13.3nM > Cl 35.9nM), illustrating how automated profiling informs halogen selection in lead optimization.

### 6.3 Subtype-Selectivity Analysis, Melatonin Receptors MT1 and MT2 (6ME3 / 6ME6)

Closely related receptor subtypes often share similar binding pockets, allowing the same drug to bind to both but with different affinities. Explaining the structural basis of these differences is a common challenge in receptor-selective drug design. The agonist 2-phenylmelatonin (2-PMT) binds both human melatonin receptor subtypes, MT1 and MT2, but with different affinities: published radioligand binding data show that it binds MT2 (Ki = 0.17 nM) with roughly 2.5-fold higher affinity than MT1 (Ki = 0.42 nM) [17, 18]. We investigated this subtype selectivity by profiling 2-PMT bound to MT1 (PDB: 6ME3) and MT2 (PDB: 6ME6) using NCI-Profiler. In both structures, the ligand is anchored by the same conserved hydrogen-bond pair: Asn162 and Gln181 in MT1 (2.9 Å each), and the equivalent Asn175 and Gln194 in MT2 (3.0 Å and 2.9 Å) [17, 18]. The interaction profiles diverge, however, around the phenyl group of 2-PMT. In MT2, NCI-Profiler detects π–π stacking with both Phe192 and Tyr294 (Figure 5, B), whereas in MT1, the equivalent residues, Phe179 and Tyr281, fall only within hydrophobic contact range and do not meet the geometric criteria for stacking (Figure 5, A). Importantly, mutagenesis data from both original studies show that the phenylalanine position is essential for binding in both receptors: mutation of this residue (F179A in MT1; F192A or F192I in MT2) abolishes binding entirely [17, 18]. Together, these observations suggest that although the phenylalanine is important for ligand binding in both subtypes, its interaction geometry differs between them, with the absence of π–π stacking in MT1 providing a plausible structural basis for its lower affinity for 2-PMT. This case illustrates how NCI-Profiler can support subtype-selectivity analysis by comparing the interaction profile of the same ligand across related receptor structures and resolving geometric differences that may not be apparent from residue identities alone.

**Figure 5.**
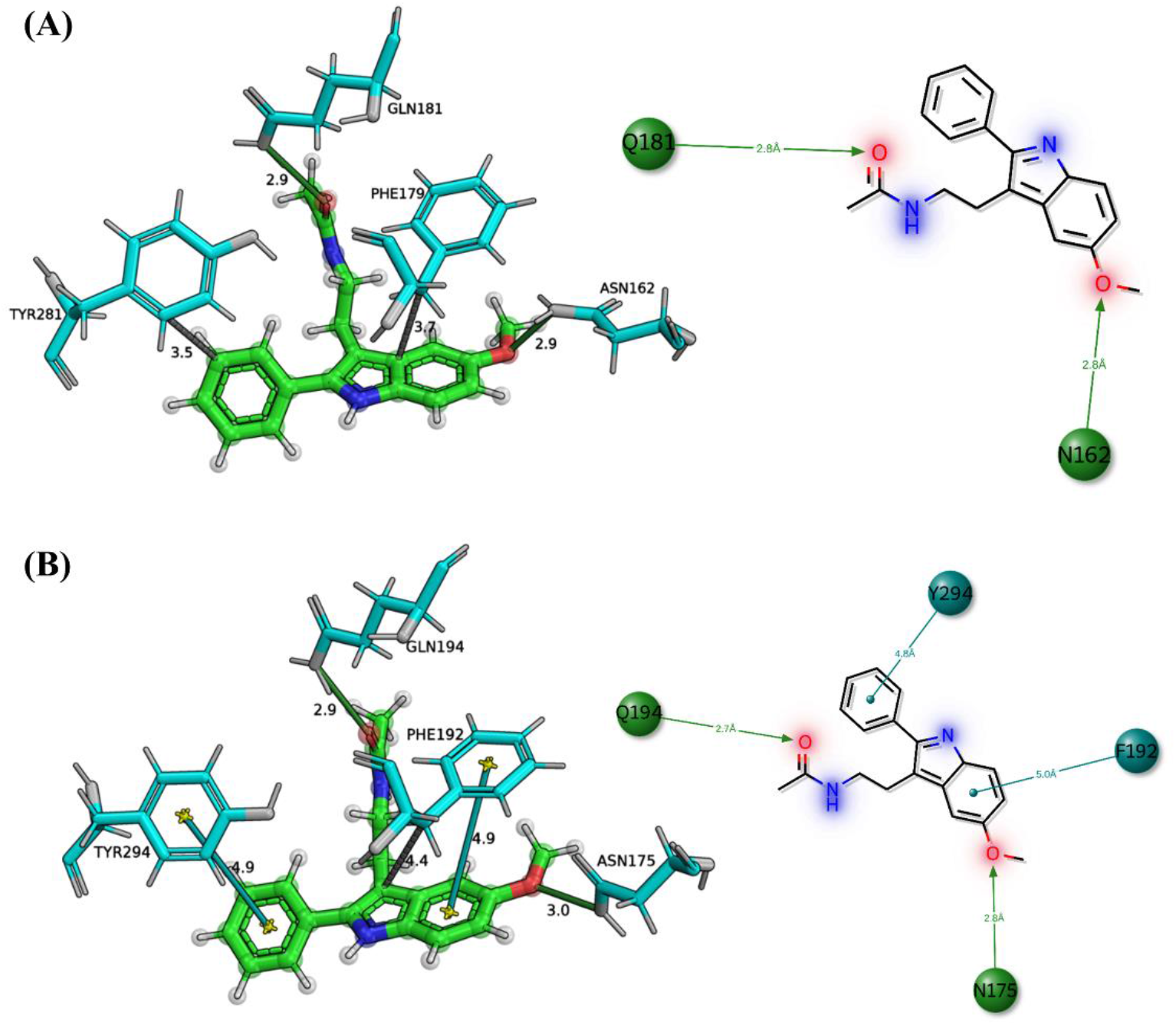
Subtype-selectivity analysis of 2-phenylmelatonin (2-PMT) bound to human melatonin receptors MT1 and MT2. NCI-Profiler interaction profiles shown as three-dimensional view (left) and 2D interaction diagram (right) for each receptor. (A) MT1 (PDB: 6ME3): 2-PMT is anchored by hydrogen bonds (green) to Asn162 (2.9 Å) and Gln181 (2.9 Å). Phe179 and Tyr281 (gray dashes, 3.7 Å and 3.5 Å) fall within hydrophobic contact range but do not meet the geometric criteria for π–π stacking. (B) MT2 (PDB: 6ME6): the same ligand forms the equivalent hydrogen bonds to Asn175 (3.0 Å) and Gln194 (2.9 Å), and additionally engages in genuine π–π stacking (teal) with both Phe192 and Tyr294 (4.8–5.0 Å). This difference in interaction geometry describes the 2.5-fold higher affinity of 2-PMT for MT2 compared to MT1.

## 7. Discussion

The validation results show that NCI-Profiler’s detection accuracy is highest for interaction types with the most geometrically constrained definitions (metal coordination, water bridges, π–π stacking) and lowest for hydrophobic contacts, where no consensus geometric definition exists in the literature. This pattern suggests that NCI-Profiler’s performance ceiling is set less by the detection algorithm itself than by how well-defined a given interaction type is in the first place. The three exemplary cases illustrate the practical consequences of this design across different stages of drug discovery: fragment growth-vector identification, where a water-mediated halogen contact pointed to a binding pocket later validated by an independently designed analog; structure–activity interpretation, where comparing halogen bond and XWH bridge profiles across a chemical series explained a potency trend that a single-structure analysis could not; and subtype-selectivity analysis, where comparing the same ligand’s interaction geometry across two related receptors identified a specific structural distinction consistent with a measured affinity difference. Together, these cases demonstrate that broadening interaction-type coverage and enabling direct, structured comparison across structures, both central design goals of NCI-Profiler, translate into genuine interpretive value rather than completeness for its own sake.

Within the broader interaction-analysis landscape, NCI-Profiler occupies a distinct position relative to established tools such as PLIP, Arpeggio, LigPlot+, and PoseEdit, each of which has shaped how interaction detection and visualization are approached, though none currently offers an interactive three-dimensional view of detected interactions directly in the browser. A direct benchmark against PLIP on the same 85-complex validation set supports this positioning, with NCI-Profiler achieving higher accuracy than PLIP and a 2.2× speed advantage (Supplementary Information). Rather than replacing these tools, NCI-Profiler is intended for cases where their scope is more limited: metalloenzyme inhibitors and metallocofactor-containing ligands, halogenated fragments and lead series where XWH bridges are pharmacologically relevant, macrocyclic compounds that challenge standard 2D depiction workflows, and batch or comparative analyses across multiple structures where a scriptable interface and structured output are required.

The present validation, while spanning all nine interaction types across 85 complexes, does not cover every binding-site geometry or metal type, and performance on structures with resolutions above 3.0 Å or unconventional metal centers may differ from the reported F1 scores. The lower hydrophobic-contact F1 score reflects this same limitation rather than a distinct detection failure: without a community-agreed geometric definition for this interaction type, the ground truth itself is less consistent than for the other eight types. Future development will extend NCI-Profiler’s detection and visualization framework to protein–protein and protein–nucleic acid interfaces and integrate its algorithm into molecular dynamics analysis pipelines and virtual screening workflows, broadening its applicability across the wider landscape of biomolecular recognition.

## Data Availability

NCI-Profiler is freely accessible without registration at https://ncip.khu.ac.kr. The source code (version 1.0.0) is available at https://github.com/rahulsharma-bio/NCI-Profiler under the GNU AGPL-3.0 license. The repository includes scripts to download all 85 validation structures and reproduce the NCI-Profiler reports. The speed benchmark details and the 85-complex validation dataset, including all PDB identifiers, binding-site definitions, literature-derived ground-truth interactions and per-structure TP/FP/FN results, are provided as Supplementary Information.

## Acknowledgements

GitHub Copilot (Microsoft, 2025) was used as a coding assistance tool for large-scale script development and compilation during this work. All geometric parameters, interaction detection thresholds, validation datasets, benchmark results, and statistical metrics were independently defined, tested, and verified by the authors. The 2D interaction diagrams and 3D PyMOL session files presented in the figures are generated directly by the NCI-Profiler tool and were finalized with minor layout edits by the authors. All text, tables, and references were drafted, verified, and revised by the authors.

## Contribution

R.S. conceived the study, designed the methodology, developed the software, and performed the validation, investigation, formal analysis, data curation and visualization. R.S. also wrote the original draft of the manuscript. E.Y.L. supervised the study, administered the project, provided resources, acquired funding, and reviewed and edited the manuscript. All authors reviewed and approved the final manuscript.

