## Supplementary Material for "NCI-Profiler: automated protein–ligand interaction detection and visualization"

*Corresponding author

### *1. Ligand Detection and Categorization*

Prior to interaction detection, NCI-Profiler classifies every molecular component in the input structure into one of six categories, standard protein residue, common ion, solvent, cofactor, glycan, or ligand, using curated lookup tables compiled from the wwPDB Chemical Component Dictionary (CCD); nucleic acid residues are separately identified and excluded from further analysis, and any component that does not match a curated entry defaults to the ligand category. Cofactor and buffer/salt molecules (e.g., glycerol, sulfate, PEG fragments) are identified from a shared lookup table but are tracked as two separate categories, since although neither participates in ligand-binding interactions of interest, distinguishing them allows their classification accuracy to be reported independently.

This classification engine was validated against 49,613 entries from the complete wwPDB CCD, achieving an overall accuracy of 99.996% (Supplementary Figure S1). Recall was 100% for standard residues (34/34), metal ions (20/20), solvent molecules (3/3), buffer/salt molecules (46/46), nucleic acid residues (37/37), and ligands (49,433/49,433). Cofactors achieved 95.0% recall (38/40), with the two unavoidable exceptions being adenosine 5′-monophosphate (AMP) and guanosine 5′-monophosphate (GMP), both of which are present in the nucleic acid lookup table as well as the cofactor table and were assigned to the nucleic acid class; NCI-Profiler resolves this ambiguity consistently through evaluation-order precedence.

### *2. Detection of Functional Groups and Atoms*

Detection of non-covalent interactions in NCI-Profiler proceeds via a two-phase, rule-based procedure. In the first phase, candidate chemical groups are enumerated separately in the protein and the ligand, with each atom or atomic group evaluated against type-specific chemical criteria compiled from published structural studies, for example donor and acceptor status for hydrogen bonds, charge center for salt bridges, or hydrophobicity for hydrophobic contacts. In the second phase, candidate pairs drawn from protein and ligand are tested against geometry constraints, distance cutoffs and, where applicable, angle restrictions, that define each interaction type, and a contact is recorded only when a candidate pair satisfies all applicable criteria simultaneously. Prior to detection, NCI-Profiler extracts all ligands contained in the structure while excluding ions, nucleic acids, modified amino acids, crystallization artifacts, and solvent components.

The first-phase chemical groups relevant to each interaction type are defined as follows. A ligand carbon atom qualifies as hydrophobic provided it carries no direct bond to a polar neighbor (N, O, S, or halogen) and is not a carbonyl carbon, while protein hydrophobic atoms are taken from a curated residue dictionary spanning nine residue types (Ala, Val, Leu, Ile, Met, Pro, Phe, Trp, and Tyr). Aromatic rings are drawn from Phe, Tyr, Trp, and His in the protein, while ligand rings of 5–7 atoms are identified by depth-first cycle enumeration and validated for planarity (maximum out-of-plane deviation ≤0.5 Å) and aromaticity (bond-length variance <0.015 Å$^2$); for tryptophan and other fused ring systems, only the ring whose centroid is closest to the interaction partner is used. Hydrogen bond donors are nitrogen or oxygen atoms that can donate a hydrogen, and acceptors are nitrogen, oxygen, or sulfur atoms with available lone pairs; protein donor/acceptor assignments come from residue-specific dictionaries, ligand assignments are based on element type and bonding environment, and halogen atoms are excluded as acceptors. Halogen bond donors are Cl, Br, or I atoms connected to carbon in the ligand, with fluorine excluded, and acceptors in the protein are proximal O, N, or S atoms.

Charged group assignment follows a similar residue- and geometry-based approach. In the protein, positive charges are assigned to the side chain nitrogens of Arg, His, and Lys, and negative charges to the carboxyl groups of Asp and Glu; in ligands, positive charges are assigned to quaternary ammonium, tertiary amine, guanidinium, and amidinium groups, negative charges to carboxylate, phosphate, and sulfonate groups, and the protonation state of ligand amines is inferred from heavy-atom neighbour geometry ($sp^3$ ~109.5° vs. $sp^2$ ~120°) when explicit hydrogens are absent. Carboxylate groups are identified by a carbon atom bonded to two oxygen atoms whose C–O distances are symmetric within 0.15 Å and both lie in the range 1.15–1.35 Å, a tolerance chosen to accommodate the bond-length uncertainty of low-resolution structures, consistent with a carboxylate moiety. Guanidinium groups are detected by the presence of a carbon center with three nitrogen neighbors, matching the connectivity of the arginine guanidinium moiety and of guanidine-containing ligands, while amidinium groups are identified by approximately equal C–N distances (difference <0.12 Å) and are further validated by confirming that neither nitrogen belongs to a closed aromatic ring, which prevents false-positive assignments to pyrimidine, imidazole, or other planar nitrogen heterocycles that share the same heavy-atom connectivity but carry no positive charge. Phosphate and sulfonate groups are assigned anionic character based on the number of bonded oxygen neighbors, phosphorus atoms bonded to three or more oxygens and sulfur atoms with three or more oxygen neighbors, an approach that allows NCI-Profiler to correctly assign charged state to all common ionic groups encountered at protein–ligand interfaces using only heavy-atom coordinates.

#### *3. Detection of Interactions*

Hydrogen bonds are detected following Jeffrey (1997); salt bridges use a wider distance cutoff than the original 4.0 Å ion-pair criterion of Barlow & Thornton (1983), following PLIP, to accommodate charge delocalization across extended functional groups and moderate-resolution structures, and carry no angular component. Hydrophobic contacts use a distance range whose upper limit follows the convention adopted by ProLIF. π–π stacking is classified using the four-parameter geometric model of Calinsky & Levy (2024): centroid distance, interplane angle, elevation angle, and vertical offset together distinguish parallel, T-shaped, offset-parallel, and intermediate stacking geometries; fused ring systems (e.g., Trp, porphyrins) are handled following PInteract, evaluating only the closest ring from each fused system to prevent over-counting. Cation–π interactions are reported for each qualifying charge–ring pair in both directions, cation–π (ligand cation to protein ring) and π–cation (ligand ring to protein cation), using the threshold established by Gallivan & Dougherty (1999). Halogen bond donor angles are constrained to ensure σ-hole alignment; fluorine is excluded as it does not form a significant σ-hole. Water bridges require a water oxygen positioned between a polar protein atom and a polar ligand atom, with a bridging angle measured at the water oxygen, and each water molecule may mediate at most two such bridging contacts. XWH bridges extend the same σ-hole criterion to water-mediated contacts: a ligand halogen engages a water molecule via σ-hole geometry, and the water simultaneously hydrogen-bonds to a protein residue using HBPLUS criteria. Metal coordination is evaluated for a broad range of biologically relevant metal ions against coordinating protein, ligand, and water atoms (N, O, S), using metal-specific distance ranges derived from crystallographic surveys, with coordination geometry determined by comparing observed donor angles against ideal angle signatures for candidate geometries across coordination numbers 2–8; when a metal bridges both protein and ligand, NCI-Profiler reports the interaction as bridged coordination. Complete distance and angle thresholds for all nine interaction types are listed in Table S1.

**Table S1.** Standard geometric thresholds used for interaction detection. Angles in degrees, distances in Ångström.

| # | Parameter | Threshold | Ref. |
|---|---|---|---|
| 1 | Hydrogen bond: D···A distance | 2.5–3.5 Å | [1] |
| | Hydrogen bond: ∠(D–H···A) | ≥ 120° | [1] |
| 2 | Salt bridge: charge-center distance | ≤ 5.5 Å | [2] |
| 3 | Hydrophobic: C···C distance | 3.3–4.5 Å | [3] |
| 4 | π–π stacking: centroid distance, d | ≤ 5.0 Å | [4] |
| | π–π stacking: interplane angle, P | ≤ 30° (parallel); ≥ 60° (T-shaped) | [4] |
| | π–π stacking: elevation angle, Tθ | ≥ 65° (true π-stacking) | [4] |
| | π–π stacking: vertical distance, h | 3.0–4.2 Å | [5] |
| 5 | Cation–π: charge–centroid distance | ≤ 6.0 Å | [6] |
| 6 | Halogen bond: X···A distance (Cl/Br/I) | 3.3 / 3.4 / 3.5 Å | [7] |
| | Halogen bond: ∠(C–X···A) | ≥ 140° | [8] |
| | Halogen bond: ∠(X···A–Y) | 90–140° | [8] |
| 7 | Water bridge: water–polar distance | ≤ 3.5 Å | [1] |
| | Water bridge: bridging angle, ω | 80–140° | [8] |
| 8 | XWH bridge: X···Ow distance (Cl/Br/I) | 3.27 / 3.37 / 3.50 Å | [9] |
| | XWH bridge: ∠(C–X···Ow) | ≥ 130° | [10] |
| | XWH bridge: Ow···D/A distance | ≤ 3.9 Å | [11] |
| 9 | Metal coordination: Zn–L distance | 1.9–2.6 Å | [12] |
| | Metal coordination: Ca–L distance | 2.2–2.8 Å | [12] |

### *4. Interaction reduction*

Raw geometric detection inevitably produces redundant contacts where the same atom or residue satisfies criteria for multiple interaction types. Reporting all such contacts inflates counts and generates profiles inconsistent with per-residue energy decomposition analyses. NCI-Profiler resolves this redundancy through an eight-step specificity-aware priority cascade operating at both atom and residue levels, following a hierarchy of decreasing geometric constraint and pharmacological distinctiveness: metal coordination > salt bridge > halogen bond > hydrogen bond > water bridge > hydrophobic contact [3]. Atoms coordinating a metal ion are first removed from the hydrogen bond, salt bridge, and halogen bond pools, preventing misclassification of coordination bonds as generic polar contacts [13, 14]. Hydrogen bond contacts involving atoms already assigned to a charged pair are then excluded from the hydrogen bond list, as are hydrogen bonds to a protein acceptor already engaged in a halogen bond, since halogen bonds are rarer and more directional [15]; among any remaining candidates, only the best-angled hydrogen bond (closest to 180°) is retained per donor hydrogen. Aromatic ring atoms already assigned to a π–π stacking pair are removed from the hydrophobic pool, water bridges are suppressed for any residue that already forms a direct polar contact (hydrogen bond, salt bridge, halogen bond, or metal coordination), and, at the residue level, any residue with a specific polar interaction has all of its hydrophobic contacts suppressed, aligning profiles with per-residue MM-PBSA decomposition granularity [16]. Finally, hydrophobic contacts are clustered so that only the shortest ligand–protein contact per residue is retained in each direction.

### *5. 2D Interaction Diagram Generation*

NCI-Profiler generates interactive 2D interaction diagrams through a multi-step pipeline that transforms the three-dimensional atomic coordinates of a protein–ligand complex into a two-dimensional schematic representation. The process consists of four stages: ligand 2D coordinate generation, interacting residue placement, interaction line rendering, and SVG assembly.

**Stage 1: Bond order and molecular assembly.** Before 2D coordinates are generated, the ligand is first assembled as a chemically correct molecular object with resolved bond orders. For known metalloporphyrins (heme, chlorophyll, and related cofactors), RDKit's direct PDB parsing is used to preserve ring connectivity, and bond orders are then corrected using a dedicated porphyrin-specific routine, since RDKit's PDB parser otherwise assigns all bonds as single. For other ligands, OpenBabel converts the PDB coordinates to SDF format, which infers proper bond orders from atomic geometry. If OpenBabel is unavailable or this step fails, NCI-Profiler falls back to standard RDKit PDB parsing.

**Stage 2: Ligand 2D coordinate generation.** For ligands containing a coordinating metal bound to three or more nitrogen atoms within 2.5 Å, characteristic of a metalloporphyrin-like ring system (e.g., heme, chlorophyll, cobalamin), NCI-Profiler first queries the PDB Chemical Component Dictionary (CCD) through PDBe CCDUtils for a template-based 2D depiction. If no CCD template is available, a PCA-based projection is used instead, in which the 3D coordinates are projected onto their best-fit plane, preserving the natural geometry of these planar ring systems; among this fallback and the standard layout algorithms below, whichever produces the fewest bond crossings is kept. For all other ligands, 2D coordinates are generated by a competitive evaluation of two layout algorithms, CoordGen and RDKit's rdDepictor, and the layout with fewer bond crossings is selected.

**Stage 3: Residue placement.** Each interacting residue label is anchored at the centroid of the ligand atom(s) it contacts and assigned to one of eight 45° angular sectors around the ligand center, based on the direction from the ligand center to that anchor point. Residues are placed outward from their anchor at a fixed minimum distance, with residues sharing a sector spread apart by an angular offset of approximately 15° and staggered radially to avoid initial overlap. An iterative collision-resolution step then checks all label pairs and pushes apart any that remain closer than a minimum separation distance, for up to twenty iterations; a label cannot be displaced more than 6.0 Å from its anchor point and remains at least 3.0 Å from it so the connecting interaction line stays visible. Residue labels are rendered as rounded rectangles, color-coded by interaction type to match the 3D PyMOL visualization, and shaded by amino acid physicochemical property; water molecules are shown as red spheres and metal ions as grey circles with element and charge labels.

**Stage 4: Interaction line rendering.** Each detected non-covalent interaction is drawn as a color-coded line connecting the relevant ligand atom to the interacting residue label, using the same color scheme as the 3D PyMOL visualization. Hydrogen bonds, salt bridges, π–π stacking, cation–π, and metal coordination are drawn as solid lines, with hydrogen bonds additionally showing a directional arrowhead pointing toward the acceptor; hydrophobic contacts, halogen bonds, water bridges, and XWH bridges are drawn as dashed lines. Distance labels (in Å) can be toggled on or off along each interaction line; for water- and metal-mediated interactions, two distance labels are shown, one on each side of the bridging atom.

**Stage 5: Interactive SVG assembly.** The completed diagram is rendered as an SVG element embedded in the web page, with draggable residue labels, a toggleable interaction-type legend, optional distance labels, and zoom controls, implemented through the web interface's JavaScript rather than embedded within the SVG file itself. For export, the diagram is rasterized from the SVG at 600 DPI and downloaded as a PNG image directly from the browser, without server-side processing.

#### *6. 3D interactive viewer*

The NCI-Profiler 3D viewer integrates Mol* v5.11.0 as a rendering component while retaining all interaction detection and interpretation within NCI-Porfiler [17]. Interaction detection, classification, filtering, and geometric definition are performed by the same NCI-Profiler analysis pipeline used for the tabulated results and 2D interaction diagrams. For 3D visualization, the resulting interactions are converted into explicit spatial data, with each interaction represented by the coordinates required to define its graphical elements. These include atom-pair coordinates for direct interactions, group or ring centers where appropriate, and the coordinates of bridging water or metal atoms for bridged interactions. Thus, Mol* receives only the structural data required for visualization together with the interaction type, display color, and associated labels; interaction criteria, thresholds, and classification decisions remain entirely within NCI-Profiler. This design ensures that the contacts displayed in 3D are the same NCI-Profiler defined interactions reported by the analysis pipeline.

The transfer from NCI-Profiler to Mol* occurs through a dedicated 3D data interface. The analyzed structure is supplied separately as the corresponding PDB file, while a server request returns the NCI-Profiler interaction data with their resolved 3D coordinates. The web client converts these data into a Mol*-compatible scene description containing the molecular structure, ligand and protein representations, interaction geometries, and labels, and submits the complete description to Mol* in a single load operation. Each interaction class is assigned an independent identifier so that its graphical elements can subsequently be controlled without reconstructing the scene or repeating the analysis. Consequently, after the initial transfer, operations such as showing or hiding interaction classes, distance labels, residue labels, and the binding-site representation act only on the already loaded Mol* scene. NCI-Profiler therefore defines what is displayed and where it is displayed, whereas Mol* is responsible for rendering and interactive manipulation of that predefined 3D representation.

### *7. Validation*

Detailed validation of NCI-Profiler on 85 literature-based protein–ligand complexes (85 unique PDB structures). Performance was quantified using precision (TP / (TP + FP)), recall (TP / (TP + FN)), and F1-score (2 × Precision × Recall / (Precision + Recall)), calculated for each validation case by comparing the set of interacting residues reported in the literature for a given interaction type against the set NCI-Profiler detects for the same complex.

**Table S2-** Literature validation of NCI-Profiler interaction detection against established benchmark structures. Binding site identifiers follow the format Ligand Name–Chain–Residue Number (e.g., BI7-A-1244)

| # | PDB ID (Res. Å) | Binding Site | Protein / Ligand | TP | FP | FN | Ref. |
|---|---|---|---|---|---|---|---|
| **1- Cation-π** | | | | | | | |
| 1 | 2JKH 1.25 | BI7-A-1244 | Factor Xa / inhibitor | CP: TRP215, TYR99, PHE174 | PC: ARG143 | | [18] |
| 2 | 2BOK 1.64 | 784-A-1244 | Factor Xa / inhibitor | CP: TRP215, TYR99, PHE174 | CP: HIS57 | — | [19] |
| 3 | 1ACJ 2.80 | THA-A-999 | AChE / tacrine | CP: TRP84, PHE330 | — | — | [20] |
| 4 | 2Y5H 1.33 | Y5H-A-1244 | Factor Xa / inhibitor 11 | CP: TRP215, TYR99, PHE174 | — | — | [21] |
| 5 | 1UV6 2.50 | CCE-C-1206 | AChBP / carbamylcholine | CP: TRP143, TYR89, TYR185, TYR192 | CP: TRP53 | — | [22] |
| 6 | 1UW6 2.20 | NCT-A-1208 | AChBP / nicotine | CP: TRP143, TYR185, TYR192 | CP: TYR89 | — | [22] |
| 7 | 4MQS 3.50 | IXO-A-501 | M2 muscarinic / iperoxo | CP: TYR104, TYR403, TYR426 | CP: TRP400, TYR430 | — | [23] |
| 8 | 3GWV 2.35 | RFX-A-801 | LeuT / R-fluoxetine | — | PC: LYS398 | CP: ARG30, TYR108, PHE253 | [24] |
| 9 | 2B4L 2.10 | BET-A-273 | ProX / glycine betaine | CP: TRP72, TRP178, TRP225 | — | — | [25] |
| **2- Halogen bonds** | | | | | | | |
| 1 | 5U00 1.41 | 7OV-A-1001 | PDE2a / iodinated inhibitor | HAL: TYR827 (I→OH) | — | — | [26] |

| | | | | | | | |
|---|---|---|---|---|---|---|---|
| 2 | 1P5E 2.22 | TBS-A-301 | CDK2 / tetrabromobenzotriazole | HAL: LEU83 (Br→O), ILE10 (Br→O) | — | HAL: GLU81 (Br→O) | [27] |
| 3 | 3SHZ 2.45 | 5CO-A-1 | PDE5 / Cl inhibitor | — | — | HAL: TYR612 (Cl→OH) | [26] |
| 4 | 3SIE 1.93 | 5BO-A-1 | PDE5 / Br inhibitor | HAL: TYR612 (Br→OH) | — | — | [26] |
| 5 | 3DN3 1.63 | IBF-A-900 | T4 lysozyme / C6F5I | HAL: MET102 (I→SD) | — | — | [28] |
| 6 | 4LBS 1.15 | 4O8-A-402 | Aldose reductase / inhibitor | HAL: THR113 (Br→OG1) | HAL: VAL47 (Cl→O) | — | [29] |
| 7 | 4OEW 2.22 | 5IO-A-1 | PDE5 / I inhibitor | — | — | HAL: TYR612 (I→OH) | [30] |
| 8 | 4X21 2.60 | 3WH-A-501 | JNK3 / iodopyrimidine | HAL: MET146 (I→SD) | — | — | [31] |
| **3- Hydrogen bond** | | | | | | | |
| 1 | 2ZC9 1.58 | 22U-H-1501 | Thrombin / inhibitor | HBO: SER214, GLY216(×2) | — | HBO: SER214 | [32] |
| 2 | 1PXP 2.30 | CK8-A-500 | CDK2 / NU6094 | HBO: LEU83(×2), ASP145 WAT: LYS33 | — | HBO: GLU81 | [33] |
| 3 | 3EHX 1.90 | BDL-A-0 | MMP / hydroxamate | HBO: ALA182 MCO: $Zn^{2+}$ | HBO: LEU181, HIS218, HIS222, HIS228, GLU219 | — | [34] |
| 4 | 2C3L 2.35 | IDZ-A-1274 | Chk1 / indazole (8.5μM) | HBO: CYS87, GLU85 | — | — | [35] |
| 5 | 2C3K 2.60 | ABO-A-1271 | Chk1 / indazole (26nM) | HBO: CYS87, GLU85, GLU55, ASN59(×2) | HBO: PHE149, CYS87 | HBO: ASP148 | [35] |
| 6 | 1OWE 1.60 | 675-A-1001 | Urokinase / naphthamidine | HBO: ASP205, GLN208, SER206 | HBO: SER206, GLY234 | — | [36] |
| 7 | 1HVR 1.80 | XK2-A-263 | HIV-1 PR / DMP-323 | HBO: ILE50A, ILE50B, ASP25A, ASP25B | HBO: ALA28A, ALA28B | — | [37] |
| 8 | 1FKB 1.70 | RAP-A-108 | FKBP12 / rapamycin | HBO: ILE56, TYR82, ASP37, GLU54, GLN53 | — | — | [38] |
| 9 | 3ERT 1.90 | OHT-A-600 | ERα / 4-hydroxytamoxifen | HBO: ARG394, GLU353 | — | — | [39] |
| 10 | 4HJO 2.75 | AQ4-A-1001 | EGFR / erlotinib | HBO: MET769 WAT: THR766 | HBO: PRO770 WAT: THR830 | — | [40] |

| | | | | | | | |
|---|---|---|---|---|---|---|---|
| 11 | 4EKL 2.00 | 0RF-A-501 | Akt1 / GDC-0068 | HBO: ALA230, GLU228 | HBO: GLU234, GLU278 | — | [41] |
| | **4- Hydrophobic Interactions** | | | | | | |
| 1 | 5C5P 1.75 | 0E0-A-1202 | TNKS2 / magic-methyl | HPI: TYR1071 | HPI: PRO1034, PHE1035, ALA1062 | HPI: TYR1050, TYR1060 | [42] |
| 2 | 2FRA 1.90 | CRV-A-999 | Cathepsin S / CRA-27934 | HPI: PHE67, VAL157, PHE205 | — | HPI: TRP26, MET68 | [43] |
| 3 | 1J4H 1.80 | SUB-A-201 | FKBP / sulfonamide | HPI: PHE46, HIS87, ILE90 | HPI: PHE36, ILE91 | HPI: TYR26, VAL55, TRP59 | [44] |
| 4 | 2ONZ 2.80 | TMJ-A-2001 | PNMT / sulfonamide | HPI: TYR40, TYR85, TYR126 | HPI: TYR35, VAL53, ALA57, PHE182 | HPI: GLY54, LEU58 | [45] |
| 5 | 3KWJ 2.80 | 23Q-A-1 | DPP-IV / aminobenzoquinolizine | HPI: TYR666 | — | HPI: TYR631, VAL656, TRP659 | [46] |
| 6 | 1M17 2.60 | AQ4-A-999 | EGFR / erlotinib | HPI: LEU764 | HPI: VAL702, LEU768, LEU820 | HPI: LYS721, THR766 | [47] |
| 7 | 2YXJ 2.20 | N3C-A-1001 | Bcl-xL / ABT-737 | HPI: PHE97, TYR101, LEU130, VAL141, PHE146 | HPI: ALA104, ALA142 | — | [48] |
| | **5- Metal coordination bonds** | | | | | | |
| 1 | 4HHB 1.74 | HEM-A-142 | Deoxyhaemoglobin / heme ($Fe^{2+}$, Type A) | MCO: HIS87–Fe–HEM | — | — | [49] |
| 2 | 2BB6 2.00 | B12-E-200 | Transcobalamin / B12 ($Co^{3+}$, Type A) | MCO: HIS175–Co–corrin | — | — | [50] |
| 3 | 1A42 2.80 | BZU-A-300 | CA-IV / brinzolamide (Zn, Type B) | MCO: HIS94, HIS96, HIS119–Zn–BZU | — | — | [51] |
| 4 | 1AZM 2.00 | AZM-A-262 | CA-I / acetazolamide (Zn, Type B) | MCO: HIS94, HIS96, HIS119–Zn–AZM | — | — | [52] |
| 5 | 3HS4 1.10 | AZM-A-701 | CA-II / acetazolamide (Zn, Type B) | MCO: HIS94, HIS96, HIS119–Zn–AZM | — | — | [53] |

| | | | | | | | |
|---|---|---|---|---|---|---|---|
| 6 | 1JK3 1.10 | BAT-A-1 | MMP-12 / batimastat (Zn, Type B) | MCO: HIS218, HIS222, HIS228–Zn–BAT | — | — | [54] |
| 7 | 1O86 2.00 | LPR-A-702 | ACE / lisinopril (Zn, Type B) | MCO: HIS383, HIS387, GLU411–Zn–LPR | — | — | [55] |
| 8 | 1T64 2.30 | TSN-A-462 | HDAC8 / trichostatin A (Zn, Type B) | MCO: ASP178, HIS180, ASP267–Zn–TSN | — | — | [56] |
| 9 | 1W22 2.20 | NHB-A-1378 | HDAC8 / CRA-19156 (Zn, Type B) | MCO: ASP178, HIS180, ASP267–Zn–NHB | — | — | [57] |
| 10 | 3OYA 2.50 | RLT-A-601 | PFV intasome / raltegravir (Mg, Type B) | MCO: ASP128, ASP185, GLU221–Mg–RLT | — | — | [58] |
| 11 | 3THH 1.50 | ABH-A-300 | Arginase I / ABH ($Co^{2+}$ binuclear, Type B) | MCO: HIS101, ASP124, HIS126, ASP128, ASP232, ASP234–Co–ABH | — | — | [59] |
| 12 | 5YXC 1.76 | CIT-A-302 | ZinT / citrate (Zn, Type B) | MCO: HIS167, HIS176, HIS178–Zn–CIT | — | — | [60] |
| 13 | 6DZQ 2.08 | HJP-A-401 | Influenza PAN / pharmacophore (Mn, Type B) | MCO: HIS41, GLU80, ASP108, GLU119, ILE120–Mn–HJP | — | — | [61] |
| | **6- *π–π* Stacking** | | | | | | |
| 1 | 1EVE 2.80 | E20-A-1 | AChE / donepezil | PST: TRP84, TRP279 | — | — | [62] |
| 2 | 1L8B 1.73 | MGP-A-1 | eIF4E / m7GpppG | PST: TRP56, TRP102 | — | — | [63] |
| 3 | 2W26 2.08 | RIV-A-1001 | Factor Xa / rivaroxaban | PST: TYR99, PHE174, TRP215 | — | — | [64] |
| 4 | 3I1Y 2.47 | 33N-A-999 | sEH / inhibitor | PST: TRP336, HIS524 | — | — | [65] |
| 5 | 4TVJ 2.10 | 09L-A-601 | PARP2 / olaparib | PST: TYR462, TYR473 | PST: HIS428 | — | [66] |
| 6 | 4BFQ 2.55 | 083-B-1 | AChBP / VUF9432 | PST: TYR186, TYR91 | — | — | [67] |
| 7 | 1N5R 2.25 | PRM-A-951 | AChE / propidium | PST: TRP286 | — | — | [68] |
| 8 | 18GS 1.90 | GDN-A-210 | GST P1-1 / DNP-GSH | PST: TYR108 | — | — | [69] |
| 9 | 1DMW 2.00 | HBI-A-700 | PheOH / dihydrobiopterin | PST: PHE254 | — | — | [70] |

| | | | | | | | |
|---|---|---|---|---|---|---|---|
| 10 | 1TLL 2.30 | FAD-A-1452 | nNOS / FAD | PST: PHE1395 | PST: TYR1175 | — | [71] |
| 11 | 1F20 1.90 | FAD-A-1501 | nNOS FAD / FAD | PST: PHE1395, TYR1197 | PST: TYR1175 | — | [72] |
| | **7- Salt bridges** | | | | | | |
| 1 | 4JRV 2.80 | KJV-A-1001 | EGFR / inhibitor | SBR: ASP831 | — | — | [73] |
| 2 | 2Z7K 1.28 | BGU-A-400 | tRNA-guanine TGTase / guanine deriv. | — | SBR: ASP102, ASP156 | — | [74] |
| 3 | 1BJI 2.00 | DPC-A-479 | Neuraminidase / zanamivir analogue | SBR: ARG371, ARG118, ARG292, GLU119, ASP151 | — | SBR: GLU227 | [75] |
| 4 | 1PPC 1.80 | MID-E-5 | Trypsin / benzamidine | SBR: ASP189 | — | — | [76] |
| 5 | 3PBL 2.89 | ETQ-A-1 | D3 receptor / eticlopride | SBR: ASP110 | — | — | [77] |
| 6 | 7Q2N 1.70 | DSM-A-201 | β-lactoglobulin / desipramine | SBR: GLU62 | — | — | [78] |
| 7 | 7Q19 1.55 | DSM-A-201 | β-lactoglobulin / desipramine (alt.) | — | — | SBR: GLU89 | [78] |
| 8 | 5MU6 1.88 | KFK-A-601 | NMT / IMP-1088 | — | — | SBR: GLN496 | [79] |
| 9 | 3ODU 2.50 | ITD-A-1500 | CXCR4 / IT1t | SBR: ASP97, GLU288 | SBR: ASP187 | — | [80] |
| | **8- Water Bridges** | | | | | | |
| 1 | 1HPX 2.00 | KNI-B-900 | HIV-1 PR / KNI-272 | WAT: ILE50A+B, ARG8, GLY27, ASP29 | — | — | [81] |
| 2 | 1Z6F 1.60 | BO9-A-401 | PBP5 / boronic acid | WAT: SER86(×2), SER110, THR214 | — | — | [82] |
| 3 | 5J1X 2.10 | AR5-A-501 | Neuropilin-1 / Arg-5 | WAT: TRP301, GLU348, THR316, PRO317 | — | — | [83] |
| 4 | 1B56 2.05 | PLM-A-136 | E-FABP / palmitate | WAT: CYS43 | — | WAT: ARG109 | [84] |
| 5 | 1EY3 2.30 | DAK-A-500 | Enoyl-CoA hydratase / DAC-CoA | WAT: TYR137 | — | — | [85] |
| 6 | 1KT4 1.46 | RTL-A-184 | RBP / retinol | WAT: PHE96 | — | — | [86] |

| | | | | | | | |
|---|---|---|---|---|---|---|---|
| 7 | 1TT2 1.85 | THP-A-501 | Staph. nuclease / THP | WAT: TYR113(×3), LEU38, ASN118, ASP21(×2), GLU43 | — | — | [87] |
| | **9- XWH bridges** | | | | | | |
| 1 | 2UY3 1.90 | H33-A-1311 | ScCTS1 / 8-chlorotheophylline | XWH: Cl→WAT2303→TYR32, PHE210 | — | — | [88] |
| 2 | 2VIV 1.72 | VG2-A-1247 | Mexiletine fragment | XWH: Cl→WAT2096→TYR94 | — | — | [89] |
| 3 | 1G3M 1.70 | PCQ-A-712 | Estrogen sulfotransferase / PCQ | XWH: Cl→WAT847→ASP22 | XWH: Cl→WAT847→LYS21 | — | [90] |
| 4 | 1G5F 1.80 | DCE-A-601 | LinB dehalogenase / dichloroethane | XWH: Cl→WAT866→ASN38 | XWH: Cl→WAT757→ASP108, TRP109; Cl→WAT866→ASP108 | — | [91] |
| 5 | 1UV5 2.80 | BRW-A-1383 | GSK-3β / bromoindirubin | XWH: Br→WAT2019→GLU97, MET101 | XWH: Br→WAT2019→ASP200, PHE201 | — | [92] |
| 6 | 1N8U 1.80 | BDD-A-411 | CSP / 12-bromo-dodecanol | XWH: Br→WAT866→ASP9, ASN61 | — | — | [93] |
| 7 | 1SN5 1.90 | T3-C-601 | Transthyretin / triiodothyronine | XWH: I→WAT3666→THR117, THR119 | XWH: I→WAT3666→ALA108, LEU110 | — | [94] |
| 8 | 3E3D 1.55 | I3C-A-130 | Lysozyme / $I_3^-$ | XWH: I→WAT1109→CYS76 | — | — | [95] |
| 9 | 2B1P 1.90 | AIZ-A-501 | JNK3 / anilinopyrazole | XWH: Cl→WAT562→LYS93, GLU111, LEU206 | — | — | [96] |
| 10 | 1WCC 2.20 | CIG-A-1299 | CDK2 / chloro-indazole | XWH: Cl→WAT2447→LYS33, ASP145 | — | — | [97] |

**Table S3.** Summary of NCI-Profiler validation results. For each of the nine interaction types, the total TP, FP, and FN counts are aggregated to compute precision, recall, and F1-score. This table provides the quantitative basis for the F1 values reported in the main text Results section.

| Interaction Type | Abbr. | # PDBs | TP | FP | FN | Precision | Recall | F1-Score |
|---|---|---|---|---|---|---|---|---|
| Cation–π | PCT | 9 | 24 | 7 | 3 | 77.4% | 88.9% | 82.8% |
| Halogen Bond | HAL | 8 | 7 | 1 | 3 | 87.5% | 70.0% | 77.8% |
| Hydrogen Bond | HBO | 11 | 33 | 15 | 3 | 68.8% | 91.7% | 78.6% |
| Hydrophobic | HPI | 7 | 17 | 14 | 14 | 54.8% | 54.8% | 54.8% |
| Metal Coord. | MCO | 13 | 13 | 0 | 0 | 100.0% | 100.0% | 100.0% |
| π–π Stacking | PST | 11 | 19 | 3 | 0 | 86.4% | 100.0% | 92.7% |
| Salt Bridge | SBR | 9 | 11 | 3 | 3 | 78.6% | 78.6% | 78.6% |
| Water Bridge | WAT | 7 | 23 | 0 | 1 | 100.0% | 95.8% | 97.9% |
| XWH Bridge | XWH | 10 | 17 | 8 | 0 | 68.0% | 100.0% | 81.0% |

**Table S4.** Summary of PLIP's accuracy on the same 85-complex benchmark set used for NCI-Profiler validation (Table S3). PLIP was evaluated against the identical literature-derived ground truth (Table S2), using its default detection settings; PLIP does not support XWH bridge detection, so its zero TP/FP reflects a genuine absence of this capability rather than a missed detection.

| Interaction Type | Abbr. | # PDBs | TP | FP | FN | Precision | Recall | F1-Score |
|---|---|---|---|---|---|---|---|---|
| Cation–π | PCT | 9 | 21 | 0 | 6 | 100.0% | 77.8% | 87.5% |
| Halogen Bond | HAL | 8 | 6 | 2 | 4 | 75.0% | 60.0% | 66.7% |
| Hydrogen Bond | HBO | 11 | 23 | 14 | 7 | 62.2% | 76.7% | 68.7% |
| Hydrophobic | HPI | 7 | 19 | 13 | 12 | 59.4% | 61.3% | 60.3% |
| Metal Coord. | MCO | 13 | 13 | 0 | 0 | 100.0% | 100.0% | 100.0% |
| π–π Stacking | PST | 11 | 19 | 3 | 0 | 86.4% | 100.0% | 92.7% |
| Salt Bridge | SBR | 9 | 11 | 2 | 3 | 84.6% | 78.6% | 81.5% |
| Water Bridge | WAT | 7 | 14 | 5 | 5 | 73.7% | 73.7% | 73.7% |
| XWH Bridge | XWH | 10 | 0 | 0 | 17 | — | — | 0.0% |

#### *8. Speed Benchmark*

To evaluate the practical speed benefit of NCI-Profiler's protonation-free detection approach, we benchmarked NCI-Profiler against PLIP on 100 protein–ligand complexes drawn from the PDBbind and BindingDB structural datasets, selected to span a wide range of structure sizes (207 to 55,125 atoms) so as to reflect realistic usage rather than a single favorable case. Both tools were run with their default settings, one structure at a time, and restricted to a single processor core so that the comparison reflected the tools' underlying efficiency rather than differences in available hardware. NCI-Profiler completed all 100 structures in a total of 43.4 seconds (mean 0.43 s, median 0.10 s per structure), compared with 94.0 seconds for PLIP (mean 0.94 s, median 0.44 s per structure), roughly 2.2-fold faster overall and more than four times faster at the median structure size (Figure S10). Both tools completed all 100 structures without error. These results are consistent with NCI-Profiler's elimination of the external protonation step required by PLIP, supporting the improved processing speed reported in the main text.

#### *References*


1. Jeffrey GA, Jeffrey GA (1997) An introduction to hydrogen bonding. Oxford university press New York
2. Barlow DJ, Thornton JM (1983) Ion-pairs in proteins. J Mol Biol 168:867–885. https://doi.org/10.1016/S0022-2836(83)80079-5
3. Bissantz C, Kuhn B, Stahl M (2010) A Medicinal Chemist's Guide to Molecular Interactions. J Med Chem 53:5061–5084. https://doi.org/10.1021/jm100112j
4. Calinsky R, Levy Y (2024) Aromatic Residues in Proteins: Re-Evaluating the Geometry and Energetics of π–π, Cation−π, and CH−π Interactions. J Phys Chem B 128:8687–8700. https://doi.org/10.1021/acs.jpcb.4c04774
5. McGaughey GB, Gagné M, Rappé AK (1998) π-Stacking Interactions. J Biol Chem 273:15458–15463. https://doi.org/10.1074/jbc.273.25.15458
6. Gallivan JP, Dougherty DA (1999) Cation-π interactions in structural biology. Proc Natl Acad Sci 96:9459–9464. https://doi.org/10.1073/pnas.96.17.9459
7. Auffinger P, Hays FA, Westhof E, Ho PS (2004) Halogen bonds in biological molecules. Proc Natl Acad Sci 101:16789–16794. https://doi.org/10.1073/pnas.0407607101

37. O'Hagan D (1994) Rational Design of Potent, Bioavailable, Nonpeptide Cyclic Ureas as HIV Protease Inhibitors. 263:

76. Bode W, Turk D, Stürzebecher J (1990) Geometry of binding of the benzamidine- and arginine-based inhibitors $N^{\alpha}$ -(2-naphthyl-sulphonyl-glycyl)- DL - *p* -amidinophenylalanyl-piperidine (NAPAP) and (2 *R* ,4 *R* )-4-methyl-1-[ $N^{\alpha}$ -(3-methyl-1,2,3,4-tetrahydro-8-quinolinesulphonyl)- L -arginyl]-2-piperidine carboxylic acid (MQPA) to human α-thrombin: X-ray crystallographic determination of the NAPAP-

trypsin complex and modeling of NAPAP-thrombin and MQPA-thrombin. Eur J Biochem 193:175–182. https://doi.org/10.1111/j.1432-1033.1990.tb19320.x

***Supplementary Figures***

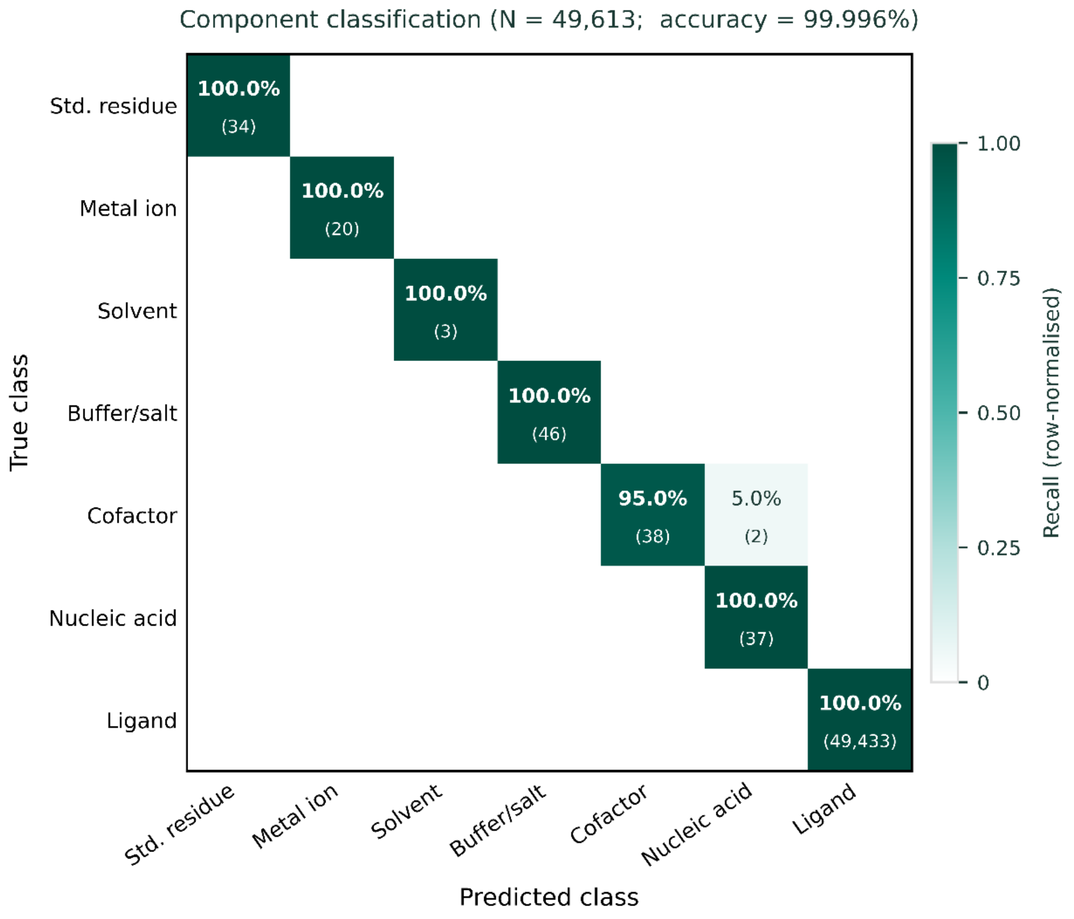


**Figure S1-** Component classification confusion matrix for NCI-Profiler, evaluated against the complete wwPDB Chemical Component Dictionary (CCD; version February 2026; N = 49,613 entries). Rows represent true classes; columns represent predicted classes. Cell color encodes per-class recall (row-normalised), scaled from white (0) to dark teal (1.0). Each non-empty cell displays the recall percentage (top) and absolute count (bottom). All seven molecular component classes standard residues, metal ions, solvents, buffer/salt molecules, cofactors, nucleic acid residues, and drug-like ligands achieved perfect or near-perfect recall. Overall accuracy: 99.996%. The two off-diagonal entries correspond to adenosine 5′-monophosphate (AMP) and guanosine 5′-monophosphate (GMP), both assigned to the Nucleic acid class instead of Cofactor owing to their presence in both lookup tables; evaluation-order precedence resolves the ambiguity consistently.

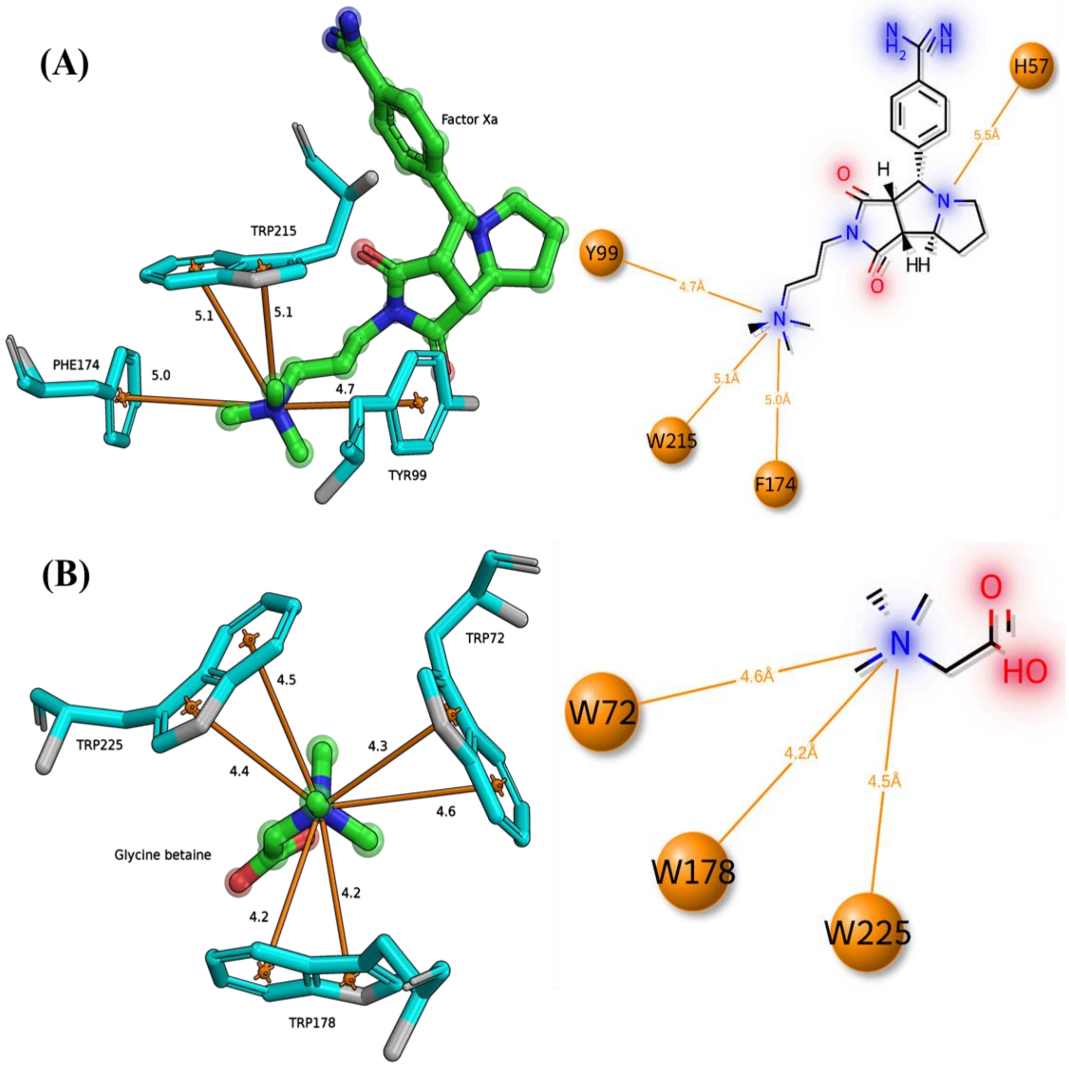


**Figure S2.** Cation–π interactions: (A) Factor Xa/784, 784-A-1244 (PDB: 2BOK); (B) ProX/glycine betaine, BET-A-273 (PDB: 2B4L). Left: PyMOL 3D view; right: NCI-Profiler 2D diagram. Cation–π contacts shown as orange lines.

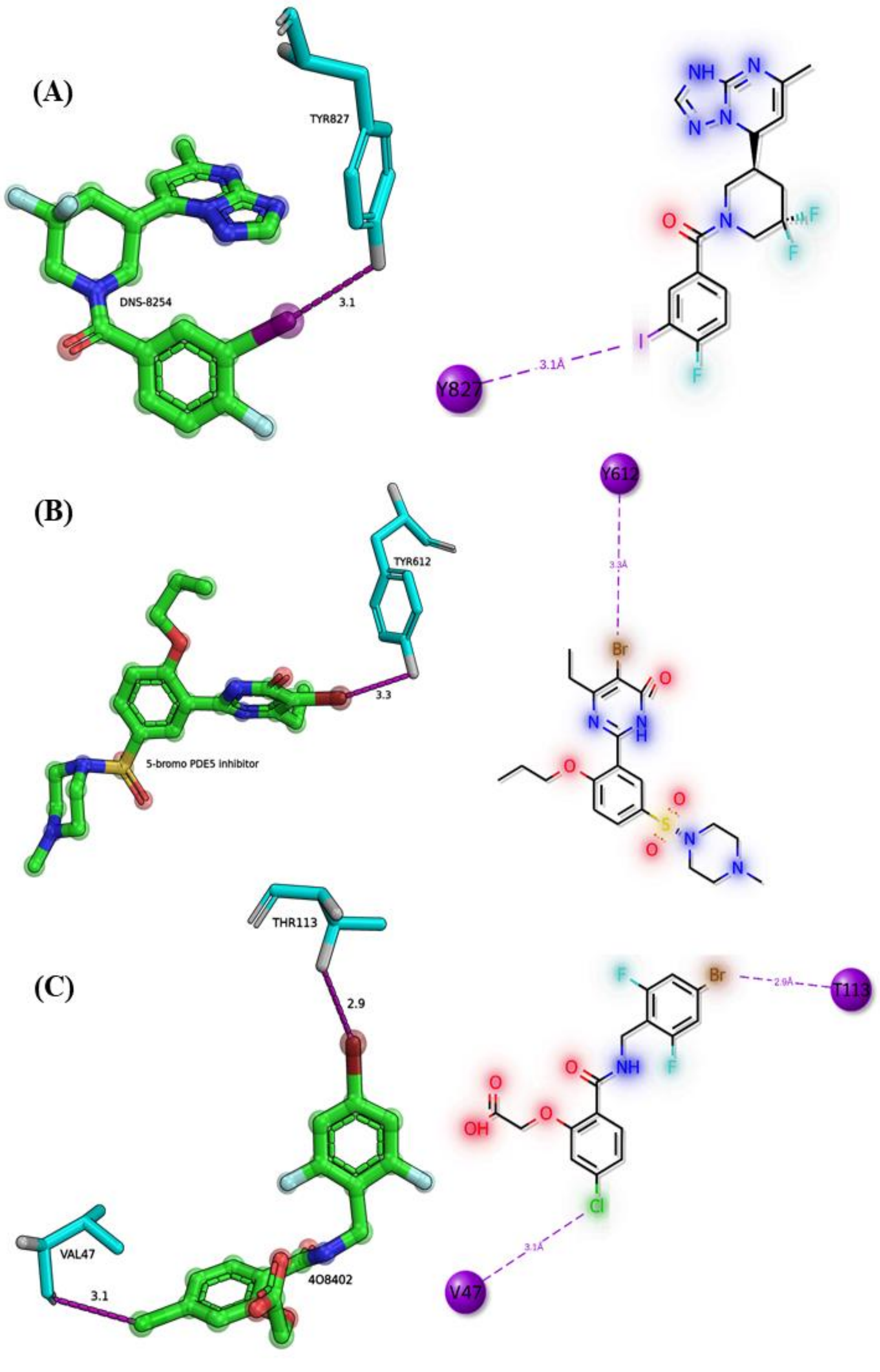


**Figure S3.** Halogen bonds: (A) PDE2a/iodinated inhibitor, 7OV-A-1001 (PDB: 5U00); (B) PDE5/Br inhibitor, 5BO-A-1 (PDB: 3SIE); (C) aldose reductase/inhibitor, 4O8-A-402 (PDB: 4LBS). Halogen bonds shown as purple dashed lines.

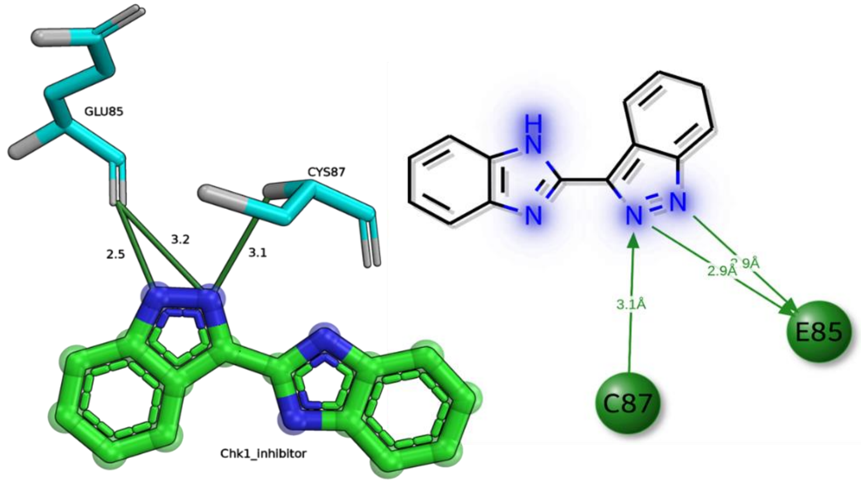


**Figure S4.** Hydrogen bonds: ERα / 4-hydroxytamoxifen, OHT-A-600 (PDB: 3ERT). Hydrogen bonds shown as green lines.

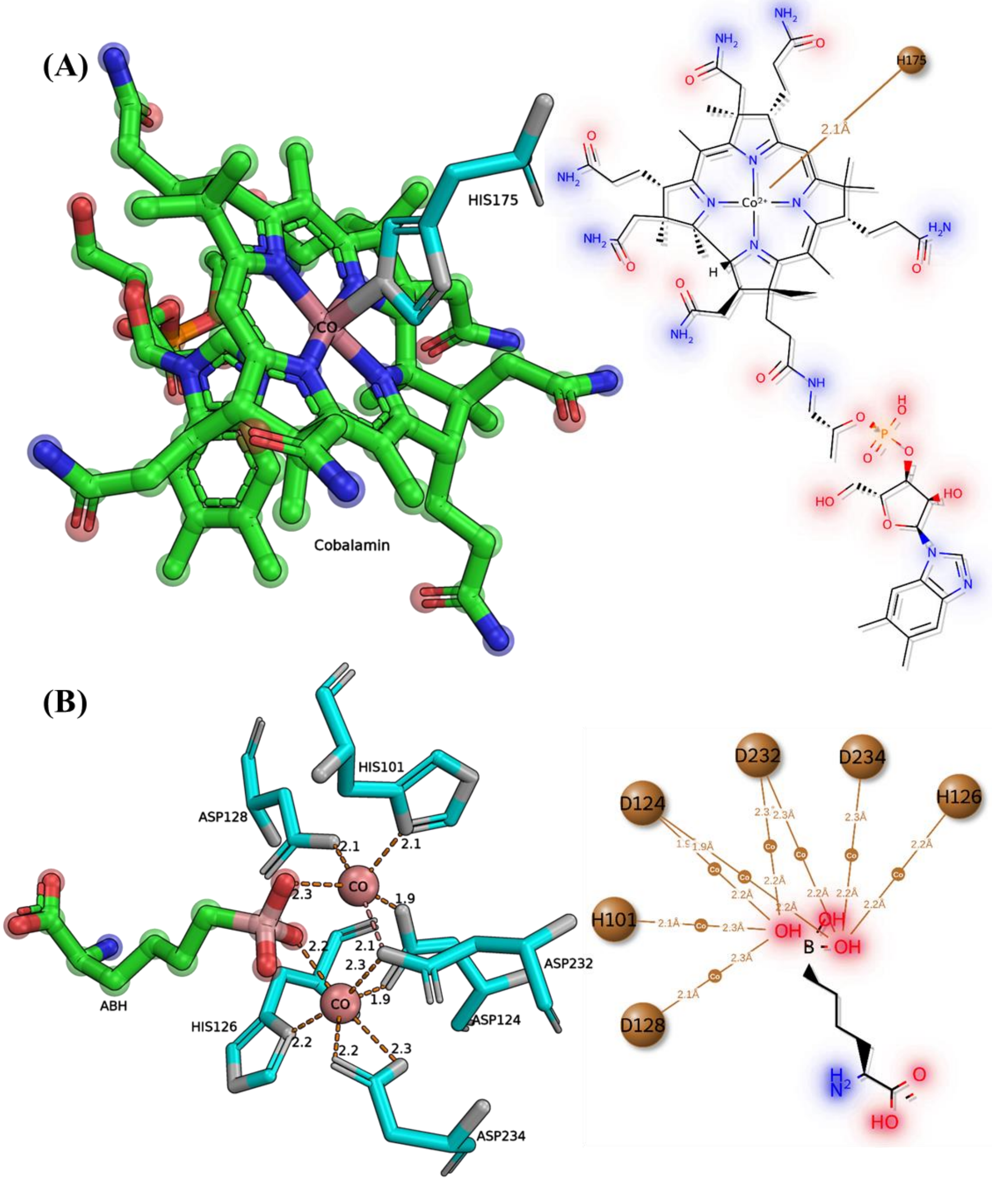


**Figure S5.** Metal coordination: (A) Type A- transcobalamin/cobalamin with $Co^{3+}$, B12-E-200 (PDB: 2BB6); (B) Type B- arginase I/ABH with binuclear $Co^{2+}$, ABH-A-300 (PDB: 3THH). Metal coordination shown as brown lines.

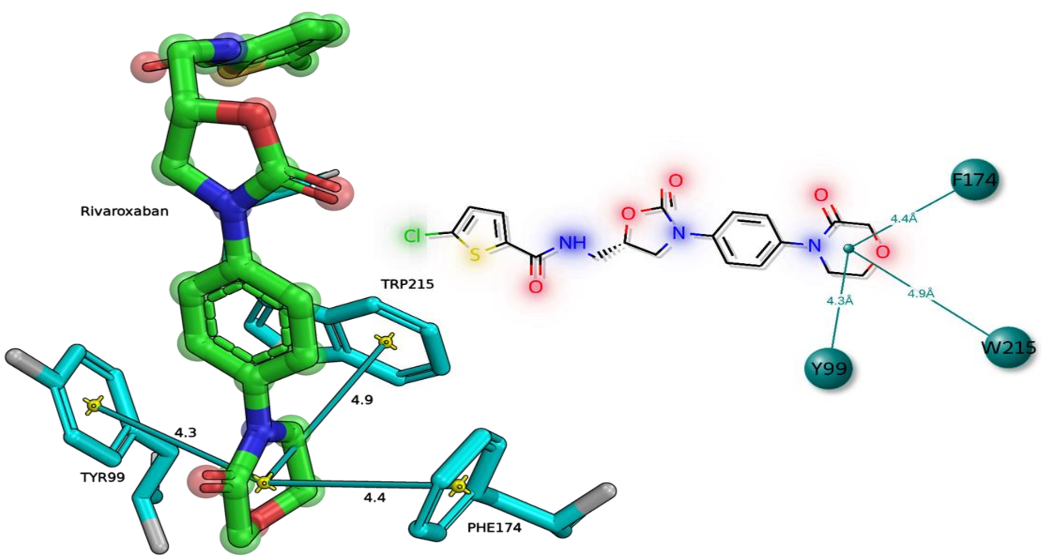


**Figure S6.** π–π stacking: Factor Xa/rivaroxaban, RIV-A-1001 (PDB: 2W26). Stacking interactions shown as teal lines.

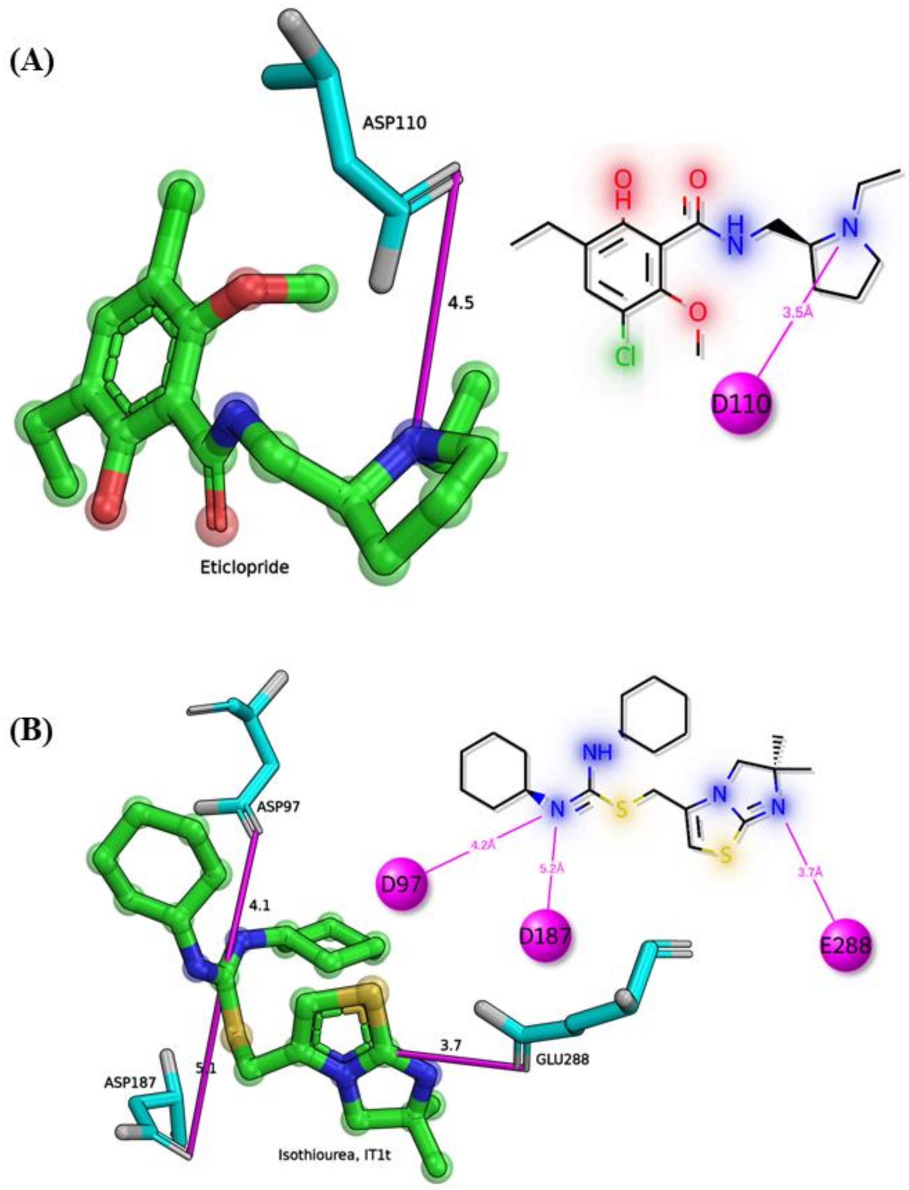


**Figure S7.** Salt bridges: (A) D3 receptor/eticlopride, ETQ-A-1 (PDB: 3PBL); (B) CXCR4/IT1t, ITD-A-1500 (PDB: 3ODU). Salt bridges shown as magenta lines.

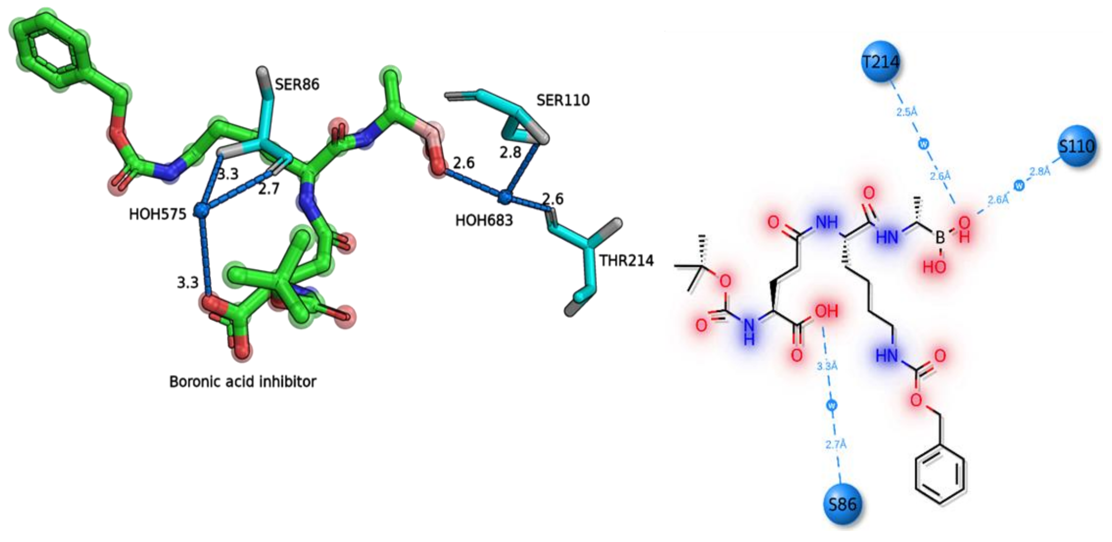


**Figure S8.** Water bridges: PBP5/boronic acid inhibitor, BO9-A-401 (PDB: 1Z6F). Water bridges shown as blue dashed lines through water intermediaries.

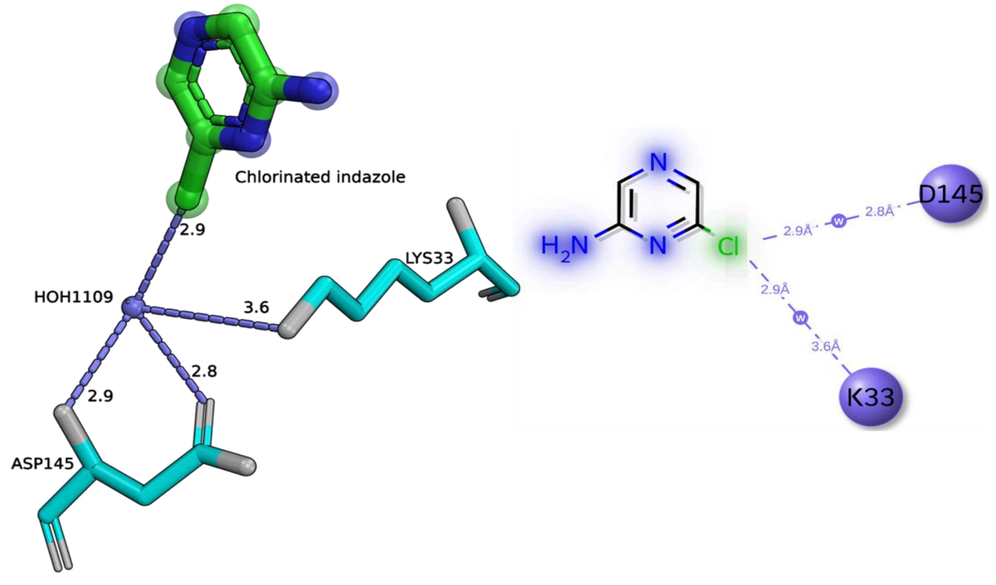


**Figure S9.** XWH bridges: CDK2 / chloro-indazole, CIG-A-1299 (PDB: 1WCC). XWH pathways shown as composite halogen→water→protein relays.

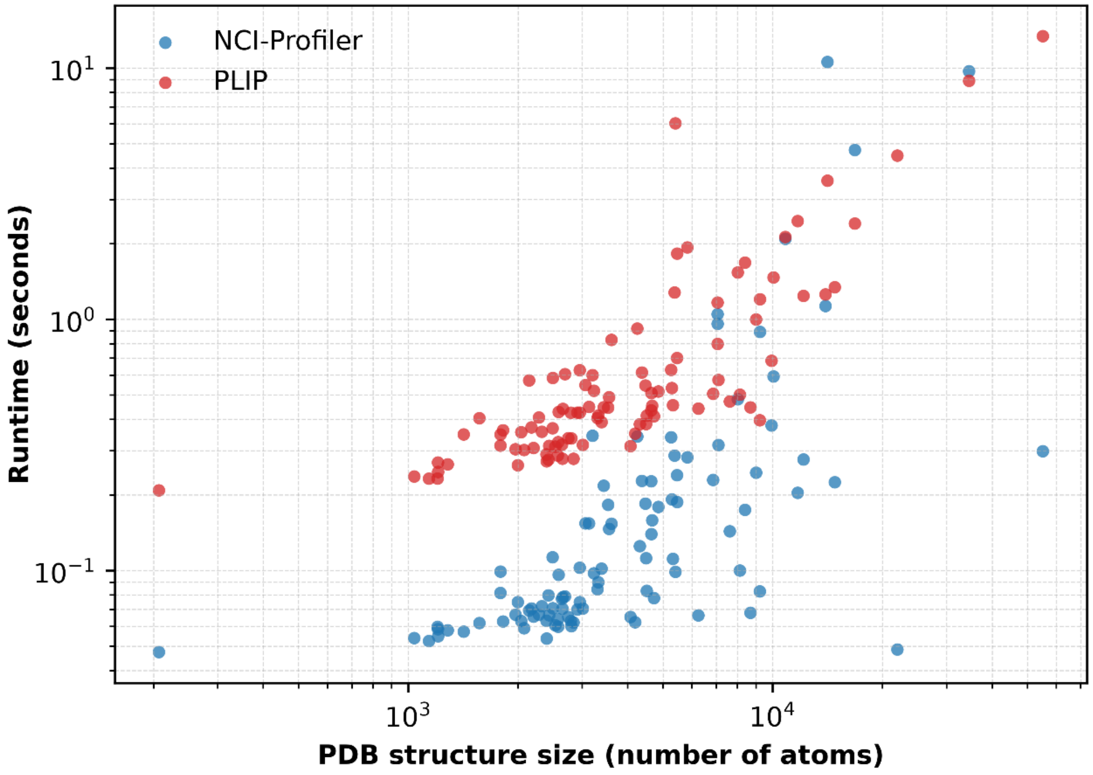


**Figure S10.** Runtime versus structure size for NCI-Profiler and PLIP across the 100-complex speed benchmark (207–55,125 atoms). Each point is one structure, run single-threaded with default settings.
